# Scenarios for the emergence of new miRNA genes in the plant *Arabidopsis halleri*

**DOI:** 10.1101/2024.07.25.605110

**Authors:** Flavia Pavan, Eléanore Lacoste, Vincent Castric, Sylvain Legrand

**Affiliations:** Univ. Lille, CNRS, UMR 8198 - Evo-Eco-Paleo, F-59000 Lille, France; Génomique Métabolique, Genoscope, Institut François Jacob, CEA, CNRS, Univ Evry, Université Paris-Saclay, 91057 Evry, France

**Keywords:** Arabidopsis, evolution, evolutionary novelty, evolution of regulatory systems, microRNA, transposable elements

## Abstract

MicroRNAs (miRNAs) are central players of the regulation of gene expression in Eukaryotes. The repertoires of miRNA genes vary drastically even among closely related species, indicating that they are evolutionarily labile. However, the processes by which they originate over the course of evolution and the nature of their progenitors across the genome remain poorly understood. Here we analyzed miRNA genes in *Arabidopsis halleri*, a plant species where we recently documented a large number of species-specific miRNA genes, likely to represent recent events of emergence. Analysis of sequence homology across the genome indicates that a diversity of sources contributes to the emergence of new miRNA genes, including inverted duplications from protein-coding genes, rearrangements of transposable element sequences and duplications of preexisting miRNA genes. Our observations indicate that the origin from protein-coding genes was less common than was previously considered. In contrast, we estimate that almost half of the new miRNA genes likely emerged from transposable elements. Miniature inverted transposable elements (MITE) seem to be particularly important contributors to new miRNA genes, with the Harbinger and Mariner transposable element superfamilies representing disproportionate sources for their emergence. We further analyzed the recent expansion of a miRNA family derived from MuDR elements, and the duplication of miRNA genes formed by two hAT transposons. Overall, our results illustrate the rapid pace at which new regulatory elements can arise from the modification of preexisting sequences in a genome, and highlight the central role of certain categories of transposable elements in this process.

## Introduction

The process by which new genes originate over the course of evolution is a driving force of evolutionary change and phenotypic innovation. A first scenario of origination is the modification of sequences that were already genes, and implies various mechanisms such as gene duplication followed by divergence, gene fusion/fission (after the deletion or insertion of a genomic region), horizontal gene transfer, or the reverse transcription and integration of mature messenger RNAs into the genome (reviewed in Van Oss and Carvunis, 2019). Gene duplication is a major mechanism in plants, as is apparent from the observation that 65% of annotated genes have a duplicate copy. Of these, most derive from whole genome duplications and/or polyploidization events, which occurred multiple times in plant evolution (Panchy et al., 2016). Yet, it is now becoming clear that novel genes can also be formed *de novo* from previously non-genic regions that have acquired gene-like characteristics. The mechanisms of *de novo* gene birth are less well understood, and have been mostly studied in the case of protein-coding genes. The *de novo* emergence of a new protein-coding gene involves the relatively complex process of acquisition of an open reading frame (ORF) along with the capacity to become transcriptionally active, although other processes can be envisioned (reviewed in Van Oss and Carvunis, 2019). The physiological relevance and evolutionary scenarios of *de novo* gene birth are still debated (Broeils et al., 2023).

MicroRNAs (miRNAs) are key regulators of gene expression in eukaryotes, and are produced from a special class of genes that do not encode for proteins, but instead are transcribed by RNA polymerase II into short RNA precursors presenting a hairpin-like structure (Bartel et al., 2004; Voinnet, 2009; Rogers and Chen, 2013). In plants, miRNAs are produced through the processing of the hairpin precursor by DICER-LIKE (DCL) proteins. Mature miRNAs are typically 21 nucleotides long and can downregulate gene expression by binding to ARGONAUTE proteins resulting in either mRNA degradation or translation inhibition (Zhan and Meyers, 2023). In plants, while some miRNA genes are deeply conserved, the majority of miRNAs are lineage-specific and thus are evolutionarily young (Fahlgren et al., 2010; Cuperus et al., 2011; Chávez Montes et al., 2014; Guo et al., 2022a; Pavan et al., 2024).

Similar to protein-coding genes, new miRNA genes can be formed from ancestral genes or have a *de novo* origin. The duplication of an existing miRNA gene can expand the family from which it originated. This process can involve whole-genome duplication, segmental duplication and tandem duplication (Sun et al., 2012). Subsequent processes of genomic diversification may follow, resulting in sub-functionalization or neo-functionalization. For example, the miR166 family in Arabidopsis expanded from whole-genome duplication, segmental duplication and tandem duplication, followed by tissue-specific subfunctionalization (Maher et al., 2006). Similarly, miR169, miR395 and miR845 families in Brassicaceae are tandemly organized and were hypothesized to originate from tandem duplications (Rathore et al., 2016). In contrast, the *de novo* origination of miRNA genes requires at least two key steps: the creation of a hairpin-like structure and the acquisition of a promoter that makes the transcription of the proto-miRNA gene possible. Three main processes have been hypothesized to allow for this dual acquisition (Nozawa et al., 2012; Cui et al., 2017; Baldrich et al., 2018). The first scenario was inspired by the observation in *Arabidopsis thaliana* of an extended similarity between the precursor sequences of some young miRNA genes and their corresponding target gene transcripts, leading to the hypothesis that they directly originate from duplications of their target genes (Allen et al., 2004). This hypothesis was later supported by specific examples in Arabidopsis (Fahlgren et al., 2010; He et al., 2014; Zhang et al., 2016), *Fragaria vesca* (Xia et al., 2015), Solanaceae (de Vries et al., 2015), Antirrhinum (Bradley et al., 2017) and Vitis (Lu et al., 2019). In this model, the inverted duplication gives rise to an initial perfectly complementary hairpin that can produce small interfering RNAs (siRNAs) also with perfect sequence complementarity to the target gene. If the duplication also involves the promoter sequence, the new miRNA gene will be expressed in the exact same tissues and stages as the gene it regulates. Over evolutionary time, the accumulation of mutations can disrupt complementarity between the two arms of the hairpin structure, facilitating recognition by the canonical DCL1 and production of miRNAs (Allen et al., 2004; Voinnet et al., 2009; Baldrich et al., 2018). While this model is particularly elegant, as it explains how new miRNA genes can gain targeting capacity right upon their inception, the proportion of new miRNA genes actually arising through this mechanism remains generally unknown.

A second source of new miRNA genes involves stem-loop sequences derived from transposable elements (TEs). This scenario of origination was supported by observations in various species including *Oryza sativa, A. thaliana* (Piriyapongsa and Jordan, 2008; Li et al., 2011; Sun et al., 2012), *Populus trichocarpa* (Sun et al., 2012), *Sorghum bicolor* (Sun et al., 2012), wheat (Poretti et al., 2019; Crescente et al., 2022) and more widely in Angiosperms (Guo et al., 2022b). TEs can be classified into class I retrotransposons, which use RNA intermediates to replicate in genomes, and class II DNA transposons, which use a “cut and paste” mechanism. Class I retrotransposons are further subdivided into long terminal repeat (LTR) retroelements such as Copia and Gypsy, and non-LTR retroelements such as LINE and SINE (Mhiri et al., 2022). Under this model, the formation of hairpin precursors can occur through the juxtaposition of two inverted LTR copies of cognate TEs. The transcription of these hairpin precursors potentially leads to the production of siRNAs, some of which may subsequently evolve into *bona fide* miRNAs (Li et al., 2011). Class II DNA transposons include various families such as Mariner, Harbinger and MuDR harboring terminal inverted repeats (TIRs) (Mhiri et al., 2022). Miniature inverted-repeat TEs (MITEs) are a special type of non-autonomous element appearing as being especially prone to generate new miRNA genes in some plant species. MITEs are truncated derivatives of autonomous Class II DNA transposons, have a short length (generally 50-800 bp long) and present terminal inverted repeats (TIRs), making them ideal substrates for the production of hairpin precursors (Guo et al., 2022b; Pegler et al., 2023). In *Oryza sativa,* up to 80% of TE-derived miRNA genes were estimated to derive from MITEs (Li et al., 2011). More recently, a study analyzed representatives from 21 species spanning from green algae to Angiosperms and suggested that 16.2% of the miRNAs loci may derive from MITEs (Guo et al., 2022b). Processing of these MITE-derived precursors by DCL proteins generates 21- or 24-nt miRNAs, which are loaded into AGO proteins and possibly modulate gene expression at either the transcriptional, *i.e.* AGO4-loaded 24-nt miRNAs, or post-transcriptional level, *i.e.* AGO1-loaded 21-nt miRNAs (Pegler et al., 2023).

A third hypothesis for the *de novo* origin of miRNA genes implies the random formation of a stem-loop structure, and the subsequent acquisition of the ability to be transcribed. Felippes et al., (2008) detected young miRNA genes in *A. thaliana* with no sequence similarity to any other region of the genome. Among them, miR823 showed homology to its orthologous intergenic region in *A. lyrata*, but this region contained two insertions, leading to a modification of the predicted secondary structure, which could explain that the homologous region in *A. lyrata* is not processed as a miRNA. Formally demonstrating the spontaneous emergence of miRNA genes from random DNA sequence motifs is difficult, and the relative contribution of the three above-described mutational processes to the influx of new miRNA genes remains poorly documented.

The Arabidopsis genus represents a unique opportunity to identify the origin of miRNA genes, by allowing the comparison of closely related genomes with a well-defined recent divergence history and solid genomic resources. The possibility to identify young miRNA genes is crucial, since they are more likely than the more anciently emerged elements to have retained nucleotide similarity to their progenitor sequence, and thus provide the “smoking guns” needed to identify the mechanisms by which they were formed. Fahlgren et al., (2010) compared the *A. thaliana* and *A. lyrata* genomes, and estimated that 85.3% of the duplications giving rise to new miRNA genes were portions of protein-coding genes. However, a limitation is that the two species diverged about 5 million years ago, which may be considered an already long time compared to the rapid dynamic of emergence of new miRNA genes. In addition, the annotation of TEs in these species has since been improved (Legrand et al., 2019), revealing a substantially higher TE content than they used, possibly leading to an underestimation of TEs as loci of origin. Finally, the miRNA annotation they used was based on a single sequencing experiment from a single accession, limiting the detection power, especially for the evolutionarily youngest miRNA genes (Pavan et al., 2024). To study more directly the process by which new miRNA genes emerge, it is thus necessary to compare species that are even more closely related. In a previous study, we identified a large number (*n* = 310) of very recent miRNA genes, specifically unique to *A. halleri* (Pavan et al., 2024), and absent from all *A. lyrata* accessions examined, from which it diverged less than one million years ago (Roux et al., 2011). In this study, we took advantage of this large set of very young miRNA genes to explore the mutational process(es) by which they emerged. Comparison of the sequences of the *A. halleri*-specific miRNA precursors with the different types of possible progenitor loci (*i.e.* protein-coding genes, TEs and other miRNA genes) revealed that transposons actually represent the most important source of origin of new miRNA genes in *A. halleri*. In particular, MITEs and certain TE superfamilies such as Harbinger and Mariner, appear to contribute preferentially. We illustrate these mechanisms of formation by documenting in detail the recent expansion of a newly created miRNA family derived from MuDR transposons, and the formation of new miRNAs by a duplication of tandemly duplicated hAT transposons.

## Results

### miRNA genes in *A. halleri* have a diversity of possible origins

The *A. halleri* reference genome (Pavan et al., 2024) comprises 34,721 predicted protein-coding genes, whose coding and non-coding fractions represent 16.7% and 20.1% of the total genome space, respectively. We annotated transposable elements using Repeatmasker and the bundle library from Legrand et al. (2019), which is composed of TEs from *A. halleri*, *A. lyrata* and *A. thaliana*. Overall, we identified 104,231 TEs sequences, comprising 31.6% of the assembly (Figure 1), in line with Legrand et al. (2019). The 463 miRNA precursor sequences predicted by Pavan et al. (2024) collectively covered 0.04% of the genome. Among them, 310 are specific to *A. halleri* accessions, as they have not been observed in any of the 13 *A. lyrata* samples examined or any other plant species (Pavan et al., 2024). These recently emerged miRNA genes may be expected to have retained a trace of their original genomic loci. To determine their progenitors, we aligned each of the 310 miRNA precursor sequences to the rest of the genome using BLASTN (Camacho et al., 2009) and cross-referenced the positions of the obtained alignments with those of protein-coding genes, transposable elements and other miRNAs.

**Figure 1:**
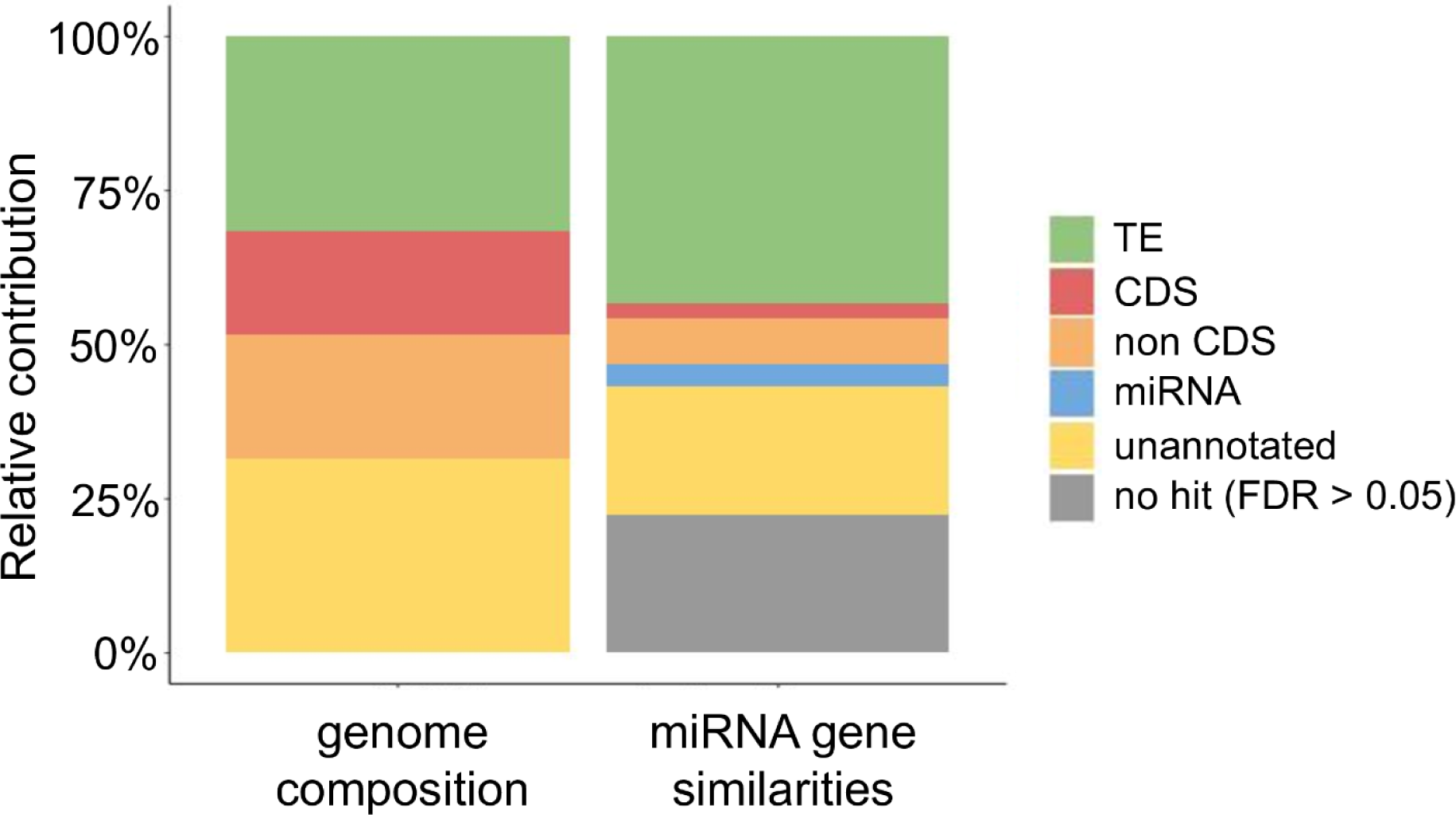
miRNA genes originate from a diversity of sources. **Left :** composition of the *A. halleri* reference genome. **Right:** relative contribution of the different sources to miRNA origination, calculated from coverage in base pairs. Each miRNA precursor sequence was compared to the *A. halleri* reference genome with BLASTN (Camacho et al. 2009) and the significant alignments (false discovery rate < 0.05) were selected. The colors represent the nature of the blast hit (existing miRNA gene, protein-coding gene, *i.e* CDS and non-CDS, and transposable element). Non-CDS annotations include introns and 5’ and 3’ untranslated regions.

We observed significant alignments for 229 of the 310 miRNA precursors, indicating that they are indeed recent enough to have preserved a trace of their origin (Figure 1). Among them, six showed similarities with CDS, 19 with intronic and/or untranslated region (UTR) sequences, 22 with other miRNA genes, 128 with TEs and 57 with unannotated regions of the assembly (Figure 1, Supplemental Table S1).

For the sake of consistency with the previous analysis by Fahlgren et al. (2010), we ensured that our results were not affected by the use of the BLASTN (Camacho et al., 2009) alignment tool rather than FASTA (Pearson, 2016), which are based on different algorithmic principles (fixed-sized words *vs.* hashing of the query) and can occasionally find matches that the other misses, and vice versa. We found qualitatively similar results using FASTA instead of BLASTN, *i.e.* the TE-related miRNAs were still a predominant source of origin (127/310) compared to protein-coding genes (46/310) (Supplementary Figure S1). Hence, we found a clear tendency for these extremely recent miRNA genes to have similarity with TE sequences (*p* < 2.2e-16), suggesting that TEs represent a major source of miRNA progenitors.

### The Harbinger, Mariner transposon superfamilies and MITEs contribute to the birth of new miRNA genes

To investigate whether particular TE sequences predominantly contribute to the origin of new miRNA genes in *A. halleri*, we compared the proportions of each TE superfamily within the set of TEs potentially associated with new miRNA origins to those in the entire annotated set of TEs in the genome assembly. In the *A. halleri* reference genome, the five most represented superfamilies are Gypsy (32.7%), Copia (19.8%), Helitron (10.9%), MuDR (10.9%) and LINE (9.2%) (Figure 2). This result is consistent with Legrand et al. (2019), who identified the same five superfamilies as the most represented in the genome of another accession (PL22). Within the set of TEs potentially associated with new miRNA origins, the most represented sequences were classified as MuDR (26.1%), MITEs (16.8%), Harbinger (9.5%) and hAT (8.1%) (Figure 2, Supplemental Table S1). In particular, while some superfamilies were underrepresented among TE-related miRNA genes such as Gypsy, Copia, SINE, and LINE (*p* < 2.2e-16), others were overrepresented such as MITE (8.5-fold higher), Harbinger (4.5-fold higher), Mariner (4-fold higher), hAT (3-fold higher) and MuDR (2.4-fold higher) (*p* < 2.2e-16), suggesting that these particular superfamilies are favored contributors to miRNA origin.

**Figure 2:**
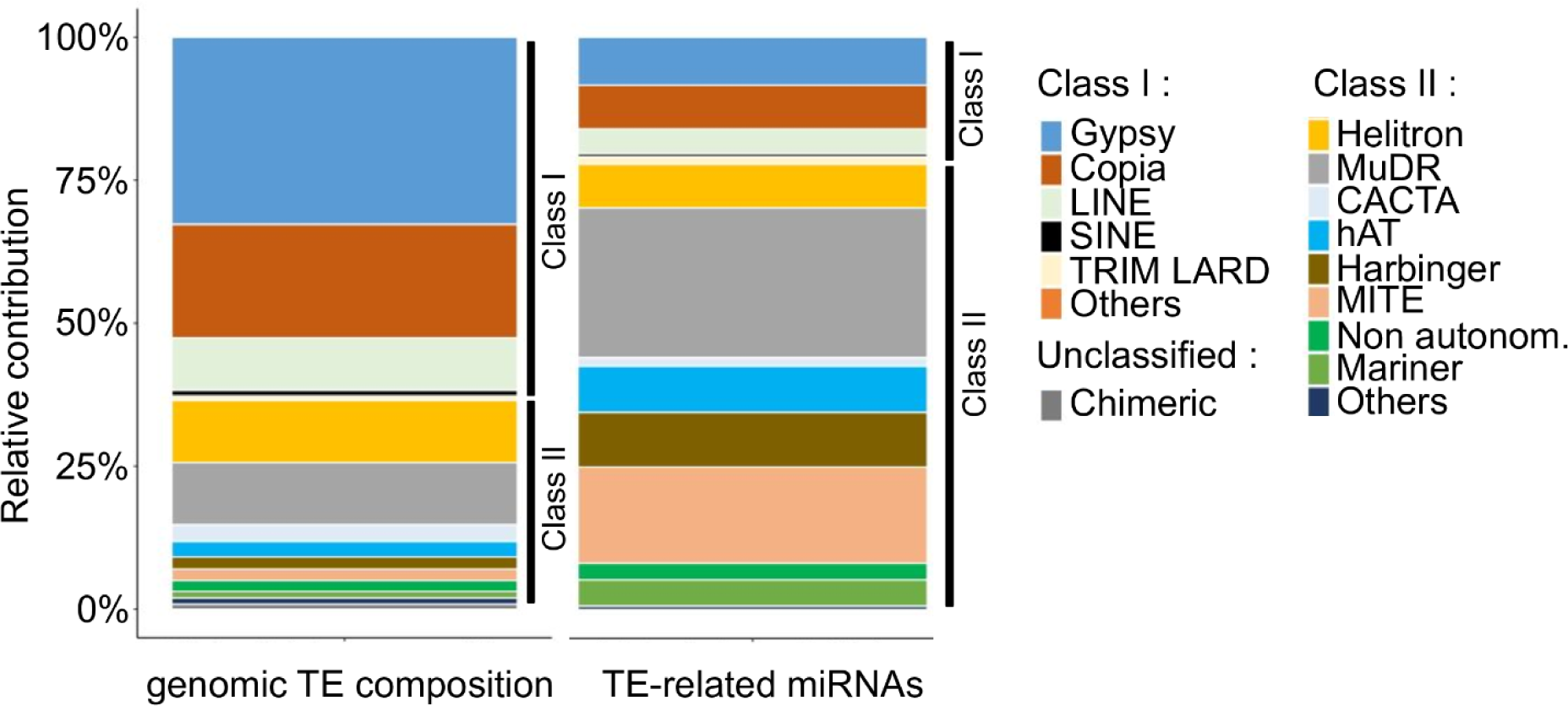
Main TE superfamilies contributing to miRNA gene birth in *A. halleri*. **Left :** relative proportions of the different TE superfamilies in the *A. halleri* reference genome. **Right:** relative proportions of the different TE superfamilies among the TE-related miRNA genes, calculated from coverage in base pairs. TE superfamilies were classified as class I retrotransposons, class II DNA transposons or unclassified transposons. Hits overlapping two or more TEs superfamilies were added to each category.

We then clustered the new miRNA genes into families based on precursor sequence similarity, and explored the distribution of the number of members (*i.e*. paralogs) in each family. The majority of the 310 new miRNA genes (*n =* 232) were classified as singletons (families of just one member). Forty miRNA genes formed clear families with two members, twenty one formed families with three members each, and a few families contained four and five members, respectively (Supplemental Figure S2). The largest family was formed by eight miRNA precursors that we named Aha-miR13a to Aha-miR13h.

### A new *A. halleri*-specific miRNA family derived from a MuDR transposon sequence

We first focused on the large eight-members family of *A. halleri*-specific miRNAs (the Aha-miR13 family). Nucleotide sequences of these miRNAs showed similarity with TEs from two different subfamilies of MuDR transposons (B-R528-Map7 and B-R2167-Map9) (Figure 3A), suggesting one, perhaps two recent expansions from this TE superfamily. Aha-miR13d, Aha-miR13f and Aha-miR13g precursors presented a large indel compared to the other members of the family. However, in spite of this relatively large indel, the secondary structure of the precursors remained largely similar (Figure 3A). Strikingly, the indel spanned the predicted miRNA-miRNA* duplex of Aha-miR13c, Aha-miR13f and Aha-miR13g (Figure 3A), but does not seem to affect the capacity of the precursor to be cleaved by DCL proteins. Most precursors produced a 21 nt-long mature miRNA sequence, suggesting that their biosynthesis is DCL1-dependent (Figure 3B). Two precursors (Aha-miR13c and Aha-miR13d) produced mature miRNA sequences with a length of 23- and 24-nt, respectively, possibly indicating that other DCL proteins may be involved in their processing (Figure 3B). In Pavan et al. (2024) we determined the preferential loading in AGO proteins of the *A. halleri* miRNAs by immunoprecipitation of AGO1 and AGO4 proteins. Interestingly, we observed that the mature miRNAs produced from the three precursors exhibiting the large indel were loaded in AGO4, while those produced by the other precursors were loaded in AGO1 (Figure 3B), suggesting a possible change of the mode of action of the family associated with the mutational event. Indeed, AGO4 generally directs DNA methylation, regulating TEs (sometimes genes) at the transcriptional level (Zhan and Meyers, 2023). Some miRNAs have been shown to be directly involved in the regulation of TEs. In *A. thaliana*, Borges et al., (2018) showed that one particular miRNA (miR845) is capable of regulating the transposon from which it is derived by cleaving TE transcripts, and initiating the production of epigenetically activated siRNAs (easiRNAs). In line with this model, we confirmed the presence of predicted targets of the AGO4-loaded 24-nt miRNAs onto the homologous MuDR sequences and other members of this superfamily. In addition, some mature miRNAs of the family are loaded in AGO1 (Pavan et al. 2024), suggesting that they could regulate mRNA targets at the post-transcriptional level. Accordingly, we observed that miRNAs produced by Ah-miR13a were predicted to be able to target the coding sequence of two protein-coding genes (Ah6g725047, related to *A. thaliana* OTU1, a deubiquitinase involved in the endoplasmic reticulum-associated degradation, Zhang et al., (2020) and Ah8g676367, homologous to a *A. lyrata* predicted protein with unknown function). Ah-miR13e was predicted to be able to target the same two genes (Ah6g725047 and Ah8g676367), plus Ah1g889505 (predicted to contain a Ribonuclease H domain related to LTR retroelements, Malik and Eickbush, 2001). Hence, members of this TE-derived miRNA family seem to have gained the capacity to target genes from the host genome in addition to target the TE superfamily from which they derive.

**Figure 3:**
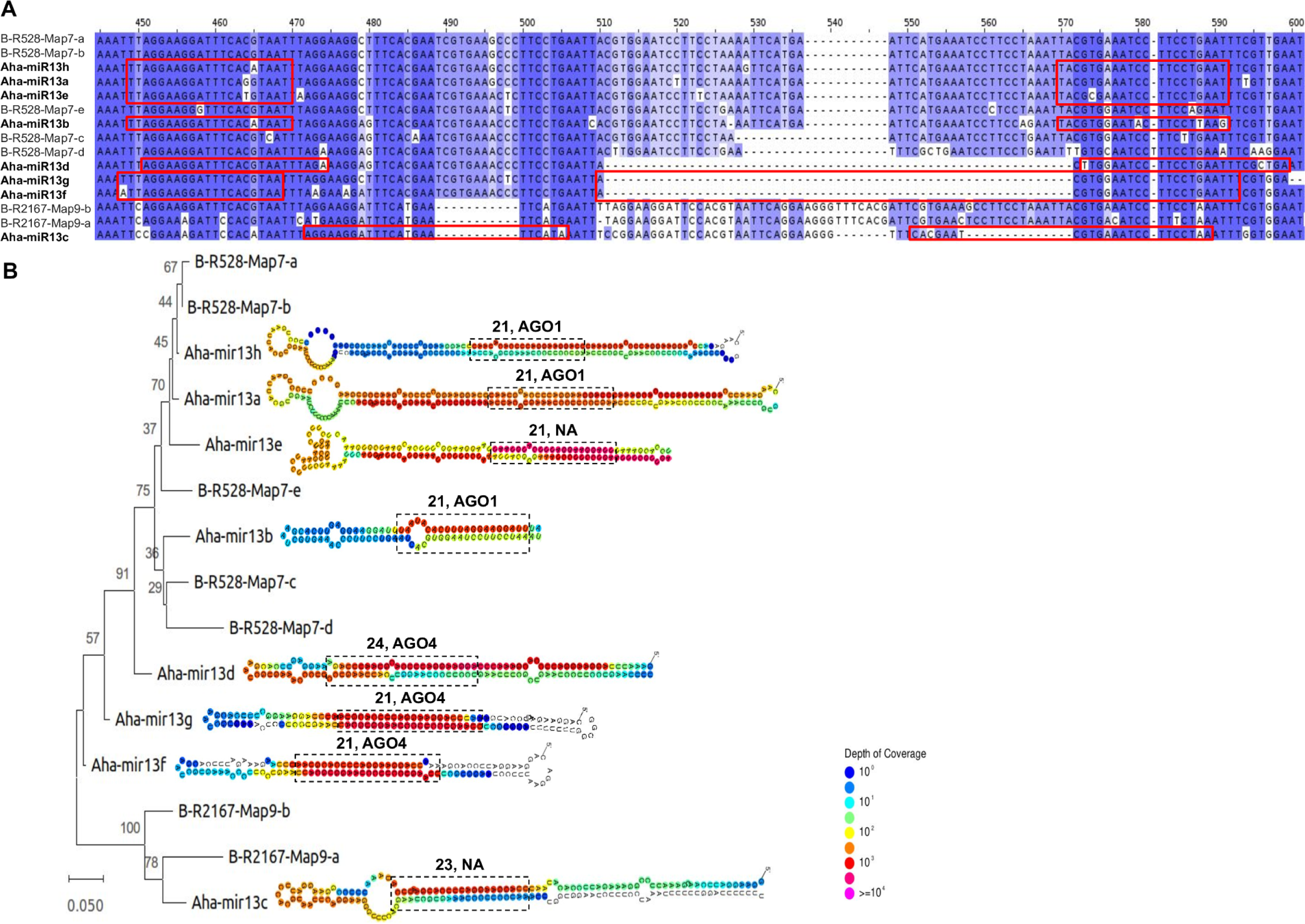
Recent expansion of a new miRNA family (Aha-miR13a-h) derived from MuDR transposon sequences. **(A)** Sequence alignment of the eight miRNA precursors and their homologous MuDR transposon sequences, aligned with MUSCLE (Edgar, 2004) and visualized with Jalview v2.11.3.2 (Waterhouse et al., 2009). Mature miRNA and miRNA* sequences are indicated by red rectangles. The color of the nucleotides represents the level of conservation, *i.e*. white for poorly conserved sites and dark blue for highly conserved sites. **(B)** Phylogenetic relationship of miRNA precursors constructed using the neighbor-joining algorithm implemented in MEGA v11.0.13 (Tamura et al., 2007). Scale bar represents genetic distance and the node numbers the bootstrap values calculated from 10,000 replicates. Alongside the phylogenetic tree, the secondary structure of miRNA precursors was generated and displayed using structVis v0.4, with a color scale indicating the read depth. The miRNA-miRNA* duplex predicted by mirkwood is indicated by a black rectangle, labeled with its size and preferential AGO protein loading (as determined in Pavan et al. 2024). “NA” indicates that the mature miRNAs were not detected in the input of AGO1 and AGO4 immunoprecipitation experiments.

### A new *A. halleri*-specific miRNA resulting from the tandem duplication of a hAT transposon

We then focused on a family (Aha-miR52) containing three paralogous miRNA precursors, Aha-miR52a, Aha-miR52b and Aha-miR52c. The sequences of these precursors overlapped with pairs of transposons from the hAT superfamily located head-to-head (Supplementary Figure S3). Aha-miR52a was shared with *A. lyrata,* in which it is present as a single copy, while Aha-miR52b and Aha-miR52c are specific to *A. halleri* (Pavan et al., 2024), suggesting that they emerged from duplication after the divergence between the two species. The secondary structure of the three miRNA precursors, notably Aha-miR52b and Aha-miR52c, appears fairly conserved, and all share the same mature 24-nt miRNA sequence. In addition, the mature miRNAs are loaded in AGO4 (Pavan et al., 2024) and have a predicted target site in the hAT sequence from which they originated, as well as in other transposons from the hAT superfamily (Figure 4A). These mature miRNAs have no target predicted in the protein-coding genes (Pavan et al., 2024). To further understand whether the miRNA precursors originated 1) from independent juxtapositions of pairs of unrelated transposons, 2) from independent tandem duplications of initially unrelated isolated transposons or 3) from the subsequent duplication of a single initial pair of transposons, we constructed a phylogenetic tree with the six transposon sequences from which Aha-miR52a, Aha-miR52b and Aha-miR52c could have derived. In scenario 1) we expect no clear phylogenetic structure. In scenario 2) we expect that the pairs of juxtaposed transposons form three independent phylogenetic clusters, while in scenario 3) we expect that members from each of the three pairs group together in the phylogenetic tree, hence forming two separate clusters. Our results showed that the two transposons forming the miRNA precursors fall into separate groups (Figure 4B), suggesting that the three members of this family represent paralogs from one single ancestral copy rather than independent juxtaposition of unrelated TEs or separate duplications of independent copies. Interestingly, the sRNAs produced by these hairpins are 24nt-long, are loaded in AGO4 and have potential targets to regulate the transposon they originated from, as well as other members of the same superfamily (Figure 4C). These observations support the possibility that TE-related miRNA genes can originate from an inverted duplication event of a transposon or from the juxtaposition of two transposons of the same family (Pegler et al., 2023).

**Figure 4:**
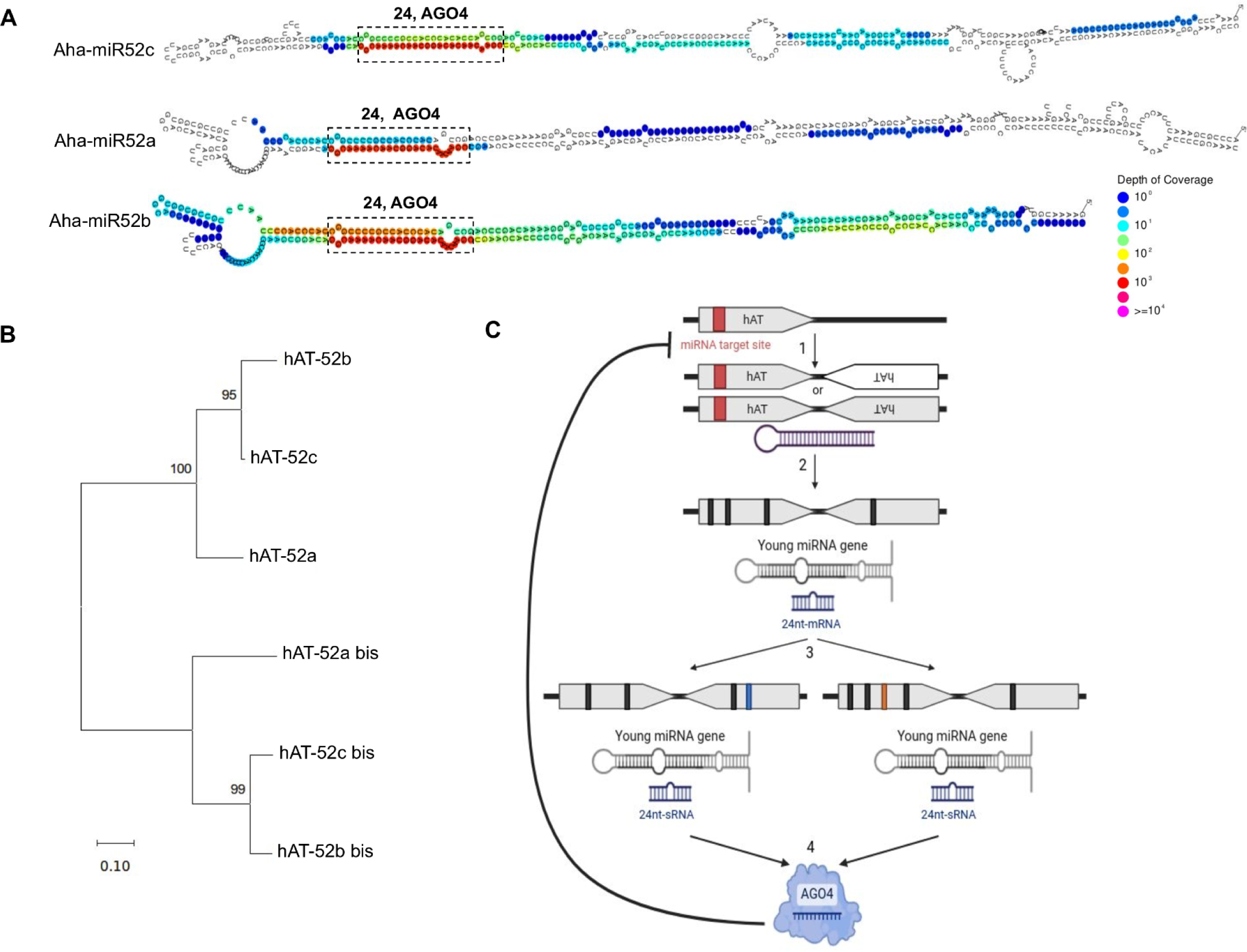
Emergence of new miRNA genes from the juxtaposition of two hAT transposons. **(A)** The secondary structures of Aha-miR52a, Aha-miR52b and Aha-miR52c precursors were obtained using structVis v0.4 and the color scale indicates the read depth. The miRNA-miRNA* duplex is indicated by a black rectangle, labeled with its size and AGO protein preferential loading. **(B)** Phylogenetic relationship of the six hAT transposons from which Aha-miR52a, Aha-miR52b and Aha-miR52c originated. The tree was constructed using MUSCLE and the neighbor-joining algorithm of MEGA v11.0.13. Scale bar represents genetic distance and bootstrap values are calculated from 10,000 replicates. **(C)** Model of emergence of new miRNA genes from hAT transposons. 1) The inverted duplication or juxtaposition of hAT transposons enables the formation of a hairpin structure. 2) Over the course of evolution, the hairpin is processed by DCL proteins and produces 24 nt-miRNAs. 3) Duplication of the region to another location in the genome expands the miRNA family. 4) The 24 nt-miRNAs produced by the hairpin are loaded in AGO4 proteins and have predicted target sites on the DNA sequence of the transposon from which they originate, as well as on those of other transposons from the same family.

### New miRNA genes can arise from the inverted duplication of a part of a coding gene

Allen et al., (2004) proposed that coding gene-derived miRNAs can arise from the reverse duplication of part of a coding gene or from a reverse intralocus duplication followed by direct duplication, and several examples of this process were reported by Fahlgren et al. (2010) in Arabidopsis. We thus looked whether some of the *A. halleri*-specific miRNA genes could have arisen this way. To illustrate this possibility, we focused on the sequence similarity we had detected between Aha-miR217 and the Ah7g813868 gene (homologous to a F-box/RNI superfamily protein-coding gene in *A. thaliana,* a large gene family with diverse roles in cell cycle transition, transcriptional regulation and signal transduction, Kuroda et al., 2002). A recent duplication of this gene giving rise to miRNA genes has been documented in *Fragaria Vesca* (Xia et al,. 2015). We compared the extended sequence of the Aha-miR217 miRNA precursor (1,000 nucleotides on each side of the loop) with the sequence of Ah7g813868. The precursor sequence was very similar to a 66 bp region in the first exon of Ah7g813868, duplicated directly, and to the same region extended to 263 bp duplicated in reverse orientation (Figure 5). This suggests that Aha-miR217 is derived from an inverted duplication of part of the Ah7g813868 gene to form a hairpin-like structure. The mature miRNA produced by the precursor (21-nt) had a predicted target site in the first exon (Figure 5), suggesting its potential to regulate its expression.

**Figure 5:**
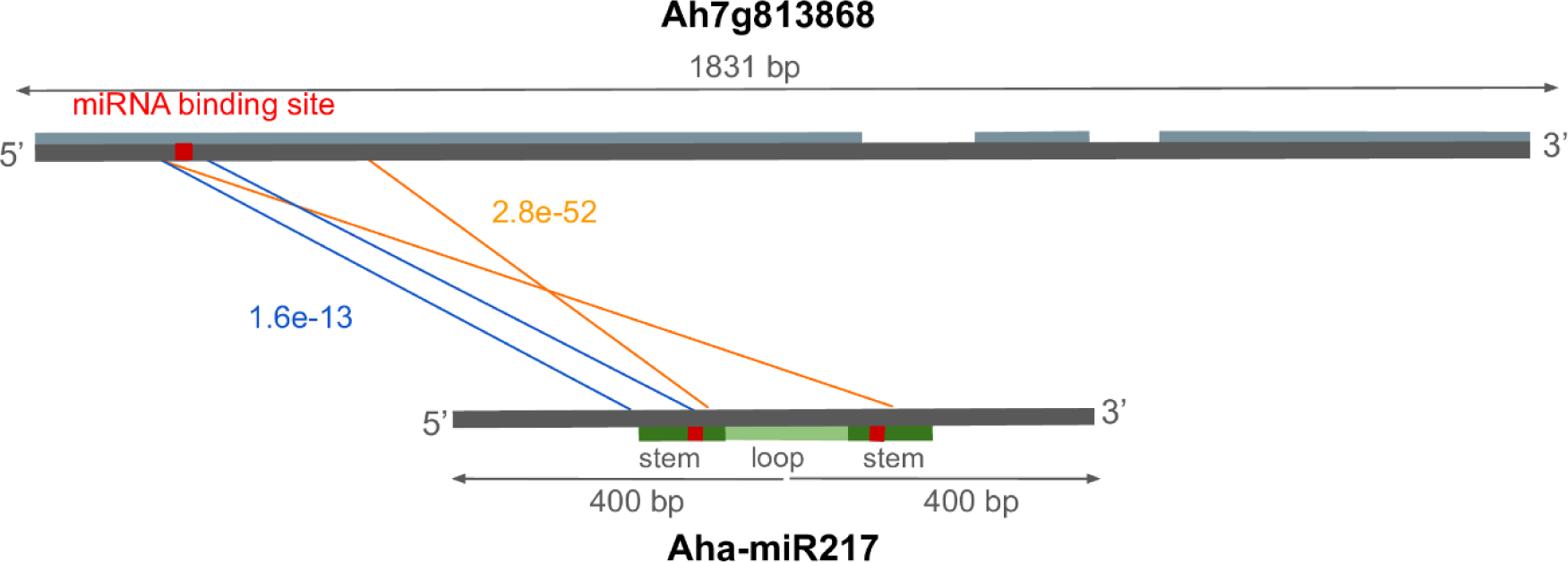
Inverted duplication of a portion of the coding sequence of a gene giving rise to a new miRNA gene. Sequence alignment between a 2000 bp region centered on the loop of the miRNA precursor Aha-miR217 and the Ah7g813868 gene using YASS (Noe and kucherov, 2005). The three exons of the gene are represented in light gray and the miRNA predicted binding site in red. The stem and the loop of the precursor are indicated in two shades of green and the miRNA and miRNA* in red. The e-value of the direct duplication is indicated in blue and the one of the inverted duplication in orange.

More generally, only six of the *A. halleri*-specific miRNA precursors have sequence similarity to CDS, and we identified inverted duplications in a very limited number of cases (only two with our criteria; Supplementary Figure S4), suggesting that this scenario of origination remains relatively rare overall.

## Discussion

### The role of transposons in the emergence of miRNA genes

Novel miRNA genes have been proposed to arise from the duplication of protein-coding genes, from transposable elements, from the duplication of preexisting miRNAs or from a region of genome able to form hairpin precursor that gain the ability to be transcribed (Nozawa et al., 2012; Cui et al., 2017; Baldrich et al., 2018). Fahlgren et al. (2010) concluded that, in *A. thaliana* and *A. lyrata,* the majority of new miRNA genes derive from protein-coding genes, with only a few with clear relation to TEs. Here, we identified a much lower fraction of miRNA genes related to protein-coding genes and instead evaluated that up to 40% of the *A. halleri*-specific miRNA genes have emerged from TEs. This difference may be due to several reasons. First, our annotation strategy was based on the comparison of a large number of distinct accessions (Pavan et al., 2024), while Fahlgren et al. (2010) predicted miRNA genes from a single sequencing experiment from a single accession. As detailed in Pavan et al., (2024), this is expected to result in a low power to detect the most recent category of miRNA genes. Second, the comparison of *A. thaliana* and *A. lyrata* is inherently limited by the relatively ancient divergence between these two species, as compared to our comparison of a pair of more closely related species, allowing for the identification of even more recent miRNA genes, providing more power to track their locus of origin. Third, TEs, due to their repetitive nature, are challenging to identify. While Fahlgren et al. (2010) used a library based on *A. thaliana* alone, our TE annotation is based on a more comprehensive library combining the repeats from *A. thaliana*, *A. lyrata* and *A. halleri* (Legrand et al., 2019). It is thus possible that Fahlgren et al. (2010) missed a number of sequence similarities between miRNAs and TEs. Fourth, the TE content in *A. thaliana* and *A. lyrata* is slightly lower than that of *A. halleri*. Legrand et al. (2019) used an unbiased estimation procedure (remapping of raw illumina reads on a joint TE library combined across the three species) and estimated the TE content in *A. thaliana* and *A. lyrata* at around 19.1% and 25.2% respectively, while in *A. halleri* it was up to 32.7% and 30.2% in the two accessions they sequenced. The estimation in our newer reference genome (31.6%) is consistent with a higher TE content in *A. halleri*, which may also contribute to explain why we identified TEs as such predominant progenitors of miRNA genes.

### Interplay between transcriptional and post-transcriptional silencing pathways

The regulation of gene expression and gene silencing can occur at two levels. Transcriptional gene silencing (TGS) allows a control of gene expression through directed methylation of the DNA sequence, while post-transcriptional gene silencing (PTGS) allows a control of gene expression through degradation of the mRNA molecules. siRNAs are main actors of TGS, while miRNAs typically function in PTGS. However, crosstalks between these two pathways have been proposed to be common and could fine-tune gene expression over the short and long terms. In Pavan et al. (2024) we showed that “young” miRNA genes tend to be produced from more “perfect” hairpins, to have a length of 24-nt, and be loaded in AGO4. Here, we show that they disproportionately stem from TE-related sequences. Collectively, these features are reminiscent of the TGS pathway. Pavan et al. (2024) further showed that, over time, the accumulation of mutations in the precursor sequence leads to the production of 21-nt miRNAs loaded in AGO1, suggesting a progressive evolution toward the PTGS pathway. In line with this scenario, we observed that the Aha-miR52 precursors potentially stemming from the duplication of hAT transposons produced 24-nt miRNAs that are loaded in AGO4 and target the transposons from which they originated. In contrast, the Aha-miR13 precursors, which probably derived from MuDR transposons, produced mainly 21-nt miRNA loaded in AGO1 and AGO4. These miRNAs target the transposon from which they originate, and in addition have gained the ability to target sites in the CDS of protein-coding genes. These two examples highlight the possible transition from the TGS to the PTGS pathways over the course of evolution: new miRNA loci predominantly derived from TE-related sequences would initially be either neutral or participate in the regulation of their progenitor TEs, as previously suggested for one particular miRNA gene from *A. thaliana* by Borges et al. (2018). The accumulation of mutations along the miRNA sequence may eventually confer the ability to target genes from the host genome. Depending on the functional importance of the newly targeted genes, such targeting may be neutral or deleterious or, under rare circumstances it may be beneficial and retained over the long run, eventually shifting to a PTGS regulatory pathway. Transitions between PTGS and TGS have been observed in other contexts. For instance, Mari-Ordonez et al. (2013) proposed that the *de novo* silencing of TEs involves an immediate PTGS response based on 21-22 nt siRNA, which is later replaced after a number of generations (eleven generations in their experiment) by a more stable, long-term TGS repression involving 24-nt sRNA molecules. Hence, TGS and PTGS mechanisms may act in concert and eventually replace each other over short and long evolutionary time scales.

### Evolution of miRNA targeting

The repertoire of targets of a miRNA gene can change over the course of evolution due to the accumulation of mutations in the miRNAs or in their mRNA target sequences. Pavan et al. (2024) observed that the number of mRNA targets tends to increase over the course of evolution, with an average of 5.4 targets for the most ancient miRNA genes, but only 0.9 for the *A. halleri*-specific miRNA genes. While TEs and protein-coding genes represent approximately similar overall fractions of the total genome, in this study, we observed that almost half of the very young miRNA genes originate from TEs, with a low proportion deriving from duplications of protein-coding genes. This observation helps to understand why the number of targets tends to increase over the course of evolution. Indeed, a miRNA gene that emerges from a protein-coding gene can immediately target the gene from which it originated. In most cases this may have deleterious consequences for the individual carrying this new miRNA gene, leading to its rapid elimination by natural selection. In contrast, a miRNA gene that emerges from a transposon is able to target the transposon from which it originates. Such a miRNA gene may be either neutral or slightly beneficial, since active transposons can invade the genome with deleterious consequences for the host. Thus, a miRNA emerging and leading to a reduction of the expression of the transposon can be retained by natural selection, at least for some time. This extension of the residence time of newly emerged miRNA genes when they arise from TEs may provide opportunity for the acquisition of new target sites in protein-coding gene through the apparition of mutations in miRNA sequence or through the insertion of the transposon of origin in a protein-coding gene leading to the creation of a new exon (exonization), a new gene or a new promoter (Etchegaray et al., 2021).

## Material and methods

### TE annotations in the *A. halleri* genome

TEs in the *A. halleri* Auby1 genome assembly were annotated using Repeatmasker (v ≥ 4.14, Smit, A.; Hubley, R.; Green, P. RepeatMasker http://www.repeatmasker.org) with the bundle library produced by Legrand et al., (2019), composed of TEs from *A. halleri*, *A. lyrata* and *A. thaliana.* Briefly, the library is composed of consensus sequences representative of TEs identified in the three species using the *TEdenovo* pipeline of the REPET package (Quesneville et al., 2005). Each consensus sequence was then classified into TE superfamilies and repeat types using PASTEC (Hoede et al., 2014).

Identification of the progenitor loci The 310 *A. halleri*-specific miRNA precursor sequences identified by Pavan et al. (2024) were aligned with BLASTN (Camacho et al. 2009) and FASTA (Pearson 2016) against the *A. halleri* reference genome assembly (Auby1 v1.3). We ignored self-alignements, and retained best-hits longer than 40 bp for further analysis. The e-values were converted to *p*-values using the relationship *p* = 1-e^-E^, and an FDR cutoff point was determined using the R (v4.1.2; R Core Team 2023) Q-VALUE package (v1.0; Storey, 2002, as in Fahlgren et al., 2010). Each significant alignment (FDR < 0.05) was intersected with protein-coding genes, TEs and miRNA gene annotations. In case the alignment overlapped with a TE within an intron, we considered that the progenitor locus was the TE. The relative proportions of the different sources of origin were normalized by base pair. Differences in the proportions of the different sources of origin between genome vs. miRNA-related loci and in the proportions of the TEs superfamilies between genome vs. miRNA-related superfamilies were tested using a χ^2^ test.

### Identification of miRNA families

We used BLASTN (Camacho et al. 2009) to compare the sequences of the miRNA precursors among them. We selected alignments with at least 80% identity, 80% coverage and an e-value < 1e-10.

Secondary structures of the miRNA precursors and their mapping density (from Pavan et al. 2024) were visualized by structVis v0.4 (github: https://github.com/MikeAxtell/strucVis).

### Phylogenetic tree

We aligned the precursor sequences with MUSCLE (Edgar, 2004) and constructed phylogenetic trees using the Neighbor Joining algorithms implemented in MEGA4 v11.0.13 (Tamura et al., 2007). Support for the nodes of the tree was evaluated with a bootstrap test (10,000 replicates).

### Target prediction

We identified the potential miRNA targets in the genome of *A. halleri* using TargetFinder (Bo and Wang, 2005) with default parameters, which provide the best balance between specificity and sensitivity (Srivastava et al., 2014). We applied a cut-off penalty score of ≤ 3 as recommended in Fahlgren et al. (2007) for reliable miRNA-mRNA target interactions.

### Origin via coding gene duplication

A region of 2kb centered on the loop of the miRNA precursor sequence was aligned to the DNA sequence of the coding gene from which it may have emerged using YASS (Noe and kucherov, 2005). Putative inverted duplications of the coding gene were inspected based on dotplots in YASS.

## Data Availability

The supplementary data set (annotation of transposable elements in *A. halleri* Auby1 reference genome) is available in Figshare under the following accession numbers : https://doi.org/10.6084/m9.figshare.26029453

## Acknowledgments

We thank the high performance computing service and Bilille at the University of Lille for providing computing resources. We thank Jean-Marc Aury, Clémentine Vitte and Hadi Quesneville for discussions and comments. We thank Idriss André and Jules Dusol (students from the MISO master) for preliminary exploration of the methods.

## Authors Contributions

Analysed data : FP and EL. Designed the project : VC, SL. Wrote the manuscript : FP, VC, SL.

## References

Allen, E., Xie, Z., Gustafson, A.M., Sung, G.-H., Spatafora, J.W., and Carrington, J.C. (2004). Evolution of microRNA genes by inverted duplication of target gene sequences in Arabidopsis thaliana. Nat Genet 36: 1282–1290.

Baldrich, P., Beric, A., and Meyers, B.C. (2018). Despacito: the slow evolutionary changes in plant microRNAs. Current Opinion in Plant Biology 42: 16–22.

Bartel DP. (2004). MicroRNAs: genomics, biogenesis, mechanism, and function. Cell:116:281–97.

Bradley, D. et al. (2017). Evolution of flower color pattern through selection on regulatory small RNAs. Science 358: 925–928.

Bo, X. and Wang, S. (2005). TargetFinder: a software for antisense oligonucleotide target site selection based on MAST and secondary structures of target mRNA. Bioinformatics 21: 1401–1402.

Borges, F., Parent, J.-S., Van Ex, F., Wolff, P., Martínez, G., Köhler, C., and Martienssen, R.A. (2018). Transposon-derived small RNAs triggered by miR845 mediate genome dosage response in Arabidopsis. Nat Genet 50: 186–192.

Broeils LA, Ruiz-Orera J, Snel B, Hubner N, and Van Heesch S. (2023) Evolution and implications of de novo genes in humans. Nat Ecol Evol. 7(6):804–815.

Camacho, C., Coulouris, G., Avagyan, V., Ma, N., Papadopoulos, J., Bealer, K., and Madden, T.L. (2009). BLAST+: architecture and applications. BMC Bioinformatics 10: 421.

Chávez Montes, R.A., Rosas-Cárdenas, D.F.F., De Paoli, E., Accerbi, M., Rymarquis, L.A., Mahalingam, G., Marsch-Martínez, N., Meyers, B.C., Green, P.J., and De Folter, S. (2014). Sample sequencing of vascular plants demonstrates widespread conservation and divergence of microRNAs. Nat Commun 5: 3722.

Crescente, J.M., Zavallo, D., Del Vas, M., Asurmendi, S., Helguera, M., Fernandez, E., and Vanzetti, L.S. (2022). Genome-wide identification of MITE-derived microRNAs and their targets in bread wheat. BMC Genomics 23: 154.

Cui, J., You, C., and Chen, X. (2017). The evolution of microRNAs in plants. Current Opinion in Plant Biology 35: 61–67.

Cuperus, J.T., Fahlgren, N., and Carrington, J.C. (2011). Evolution and Functional Diversification of *MIRNA* Genes. Plant Cell 23: 431–442.

De Vries, S., Kloesges, T., and Rose, L.E. (2015). Evolutionarily Dynamic, but Robust, Targeting of Resistance Genes by the miR482/2118 Gene Family in the Solanaceae. Genome Biol Evol 7: 3307–3321.

Edgar, R.C. (2004). MUSCLE: multiple sequence alignment with high accuracy and high throughput. Nucleic Acids Research 32: 1792–1797.

Etchegaray, E., Naville, M., Volff, J.-N., and Haftek-Terreau, Z. (2021). Transposable element-derived sequences in vertebrate development. Mobile DNA 12: 1.

Fahlgren N, Howell MD, Kasschau KD, Chapman EJ, Sullivan CM, Cumbie JS, Givan SA, Law TF, Grant SR, Dangl JL, et al. High-Throughput Sequencing of Arabidopsis microRNAs: Evidence for Frequent Birth and Death of MIRNA Genes. PLoS ONE. 2007:2(2):e219.

Fahlgren, N., Jogdeo, S., Kasschau, K.D., Sullivan, C.M., Chapman, E.J., Laubinger, S., Smith, L.M., Dasenko, M., Givan, S.A., Weigel, D., and Carrington, J.C. (2010). MicroRNA Gene Evolution in *Arabidopsis lyrata* and *Arabidopsis thaliana*. The Plant Cell 22: 1074–1089.

Fenselau De Felippes, F., Schneeberger, K., Dezulian, T., Huson, D.H., and Weigel, D. (2008). Evolution of *Arabidopsis thaliana* microRNAs from random sequences. RNA 14: 2455–2459.

Guo, Z., Kuang, Z., Deng, Y., Li, L., and Yang, X. (2022a). Identification of Species-Specific MicroRNAs Provides Insights into Dynamic Evolution of MicroRNAs in Plants. IJMS 23: 14273.

Guo, Z., Kuang, Z., Tao, Y., Wang, H., Wan, W., Hao, C., Shen, F., Yang, X., Li. L. (2022b). Miniature Inverted-repeat Transposable Elements Drive Rapid MicroRNA Diversification in Angiosperms. Molecular Biology and Evolution 39:msac224.

He, H., Liang, G., Li, Y., Wang, F., and Yu, D. (2014). Two Young MicroRNAs Originating from Target Duplication Mediate Nitrogen Starvation Adaptation via Regulation of Glucosinolate Synthesis in *Arabidopsis thaliana*. Plant Physiol. 164: 853–865.

Hoede, C., Arnoux, S., Moisset, M., Chaumier, T., Inizan, O., Jamilloux, V., and Quesneville, H. (2014). PASTEC: An Automatic Transposable Element Classification Tool. PLoS ONE 9: e91929.

Kuroda, H., Takahashi, N., Shimada, H., Seki, M., Shinozaki, K., and Matsui, M. (2002). Classification and Expression Analysis of Arabidopsis F-Box-Containing Protein Genes. Plant and Cell Physiology 43: 1073–1085.

Legrand, S. et al. (2019). Differential retention of transposable element-derived sequences in outcrossing Arabidopsis genomes. Mobile DNA 10: 30.

Li, Y., Li, C., Xia, J., and Jin, Y. (2011). Domestication of Transposable Elements into MicroRNA Genes in Plants. PLoS ONE 6: e19212.

Lu, S. (2019). *De novo* origination of *MIRNAs* through generation of short inverted repeats in target genes. RNA Biology 16: 846–859.

Maher, C., Stein, L., and Ware, D. (2006). Evolution of *Arabidopsis* microRNA families through duplication events. Genome Res. 16: 510–519.

Malik, H.S. and Eickbush, T.H. (2001). Phylogenetic Analysis of Ribonuclease H Domains Suggests a Late, Chimeric Origin of LTR Retrotransposable Elements and Retroviruses. Genome Res. 11: 1187–1197.

Marí-Ordóñez, A., Marchais, A., Etcheverry, M., Martin, A., Colot, V., and Voinnet, O. (2013). Reconstructing de novo silencing of an active plant retrotransposon. Nat Genet 45: 1029–1039.

Mhiri, C., Borges, F., and Grandbastien, M.-A. (2022). Specificities and Dynamics of Transposable Elements in Land Plants. Biology 11: 488.

Noe, L. and Kucherov, G. (2005). YASS: enhancing the sensitivity of DNA similarity search. Nucleic Acids Research 33: W540–W543.

Nozawa, M., Miura, S., and Nei, M. (2012). Origins and Evolution of MicroRNA Genes in Plant Species. Genome Biology and Evolution 4: 230–239.

Panchy, N., Lehti-Shiu, M., and Shiu, S.-H. (2016). Evolution of Gene Duplication in Plants. Plant Physiol. 171: 2294–2316.

Pavan F., Azevedo Favory J., Lacoste E., Beaumont C., Louis F., Blassiau C., Cruaud C., Labadie K., Gallina S., Genete M., Kumar V., Kramer U., A. Batista R., Patiou C., Debacker L., Ponitzki C., Houzé E., Durand E., Aury J.M., Castric V., Legrand S. (2024). The evolutionary history and functional specialization of microRNA genes in *Arabidopsis halleri* and *A. lyrata*. bioRxiv 2024.05.03.592357; doi: 10.1101/2024.05.03.592357

Pearson WR. (2016) Finding Protein and Nucleotide Similarities with FASTA. CP in Bioinformatics. 2016:53(1).

Pegler, J.L., Oultram, J.M.J., Mann, C.W.G., Carroll, B.J., Grof, C.P.L., and Eamens, A.L. (2023). Miniature Inverted-Repeat Transposable Elements: Small DNA Transposons That Have Contributed to Plant MICRORNA Gene Evolution. Plants 12: 1101.

Piriyapongsa, J. and Jordan, I.K. (2008). Dual coding of siRNAs and miRNAs by plant transposable elements. RNA 14: 814–821.

Poretti, M., Praz, C.R., Meile, L., Kälin, C., Schaefer, L.K., Schläfli, M., Widrig, V., Sanchez-Vallet, A., Wicker, T., and Bourras, S. (2020). Domestication of High-Copy Transposons Underlays the Wheat Small RNA Response to an Obligate Pathogen. Molecular Biology and Evolution 37: 839–848.

Quesneville, H., Bergman, C.M., Andrieu, O., Autard, D., Nouaud, D., Ashburner, M., and Anxolabehere, D. (2005). Combined Evidence Annotation of Transposable Elements in Genome Sequences. PLoS Comp Biol 1: e22.

Rathore, P., Geeta, R., and Das, S. (2016). Microsynteny and phylogenetic analysis of tandemly organised miRNA families across five members of Brassicaceae reveals complex retention and loss history. Plant Science 247: 35–48.

Robinson, J.T., Thorvaldsdóttir, H., Winckler, W., Guttman, M., Lander, E. S., Getz, G., Mesirov, J.P. (2011). Integrative Genomics Viewer. Nature Biotechnology 29: 24–26.

Rogers, K. and Chen, X. (2013). Biogenesis, Turnover, and Mode of Action of Plant MicroRNAs. The Plant Cell 25: 2383–2399.

Roux, C., Castric, V., Pauwels, M., Wright, S.I., Saumitou-Laprade, P., and Vekemans, X. (2011). Does Speciation between Arabidopsis halleri and Arabidopsis lyrata Coincide with Major Changes in a Molecular Target of Adaptation? PLoS ONE 6: e26872.

Shannon, P., Markiel, A., Ozier, O., Baliga, N.S., Wang, J.T., Ramage, D., Amin, N., Schwikowski, B., and Ideker, T. (2003). Cytoscape: A Software Environment for Integrated Models of Biomolecular Interaction Networks. Genome Res. 13: 2498–2504.

Srivastava, P.K., Moturu, T.R., Pandey, P., Baldwin, I.T., and Pandey, S.P. (2014). A comparison of performance of plant miRNA target prediction tools and the characterization of features for genome-wide target prediction. BMC Genomics 15: 348.

Storey, J.D. (2002). A direct approach to false discovery rates. J. R. Stat. Soc., B 64: 479– 498.

Sun, J., Zhou, M., Mao, Z., and Li, C. (2012). Characterization and Evolution of microRNA Genes Derived from Repetitive Elements and Duplication Events in Plants. PLoS ONE 7: e34092.

Tamura, K., Dudley, J., Nei, M., & Kumar, S. (2007). MEGA4: molecular evolutionary genetics analysis (MEGA) software version 4.0. Molecular Biology and Evolution, 24(8), 1596–1599.

Van Oss, S.B. and Carvunis, A.-R. (2019). De novo gene birth. PLoS Genet 15: e1008160.

Voinnet, O. (2009). Origin, Biogenesis, and Activity of Plant MicroRNAs. Cell 136: 669–687.

Waterhouse, A.M., Procter, J.B., Martin, D.M.A, Clamp, M., Barton, G.J (2009), “Jalview version 2: A Multiple Sequence Alignment and Analysis Workbench,” Bioinformatics 25 (9) 1189–1191

Xia, R., Ye, S., Liu, Z., Meyers, B.C., and Liu, Z. (2015). Novel and Recently Evolved MicroRNA Clusters Regulate Expansive *F-BOX* Gene Networks through Phased Small Interfering RNAs in Wild Diploid Strawberry. Plant Physiol. 169: 594–610.

Zang, Y., Gong, Y., Wang, Q., Guo, H., and Xiao, W. (2020). Arabidopsis OTU 1, a linkage- specific deubiquitinase, is required for ENDOPLASMIC RETICULUM -associated protein degradation. The Plant Journal 101: 141–155.

Zhan, J. and Meyers, B.C. (2023). Plant Small RNAs: Their Biogenesis, Regulatory Roles, and Functions. Annu. Rev. Plant Biol. 74: 21–51.

Zhang, Y., Xia, R., Kuang, H., and Meyers, B.C. (2016). The Diversification of Plant *NBS-LRR* Defense Genes Directs the Evolution of MicroRNAs That Target Them. Mol Biol Evol 33: 2692–2705.

