## Supplemental data for "Scenarios for the emergence of new miRNA genes in the plant *Arabidopsis halleri*"

**Supplementary figures**

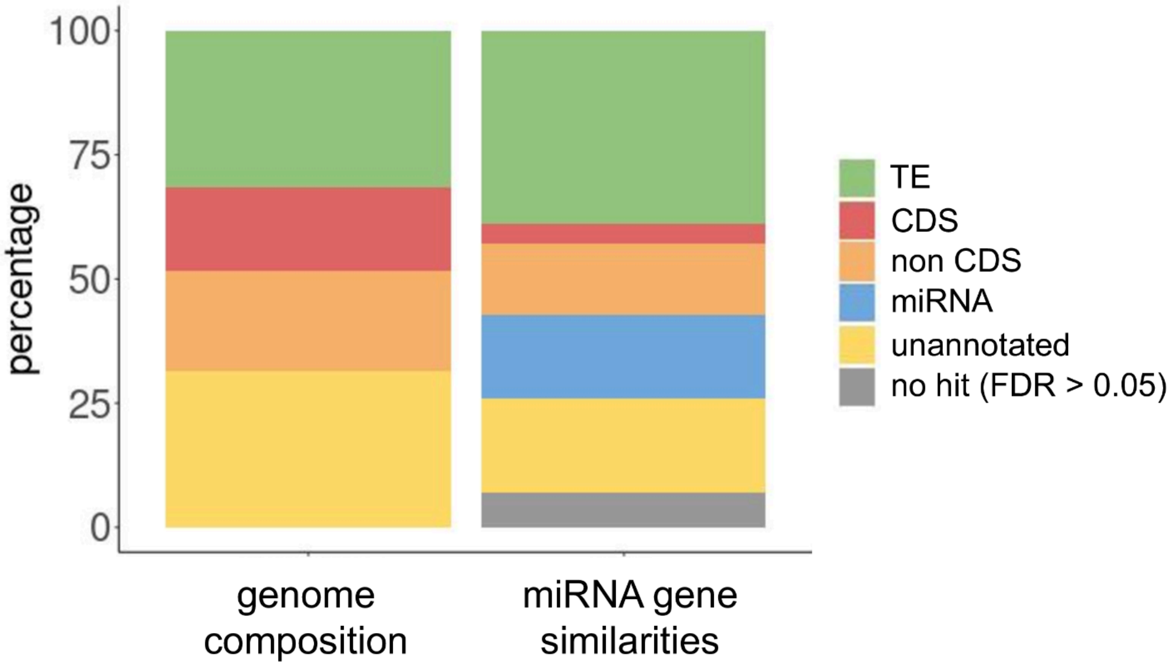


**Figure S1: miRNA genes sources of origin using FASTA.** Composition of the *A. halleri* reference genome (left) and relative contribution of the different sources to miRNA origination (right), calculated from coverage in base pairs. Each miRNA precursor sequence was compared to the *A. halleri* reference genome with FASTA (Pearson, 2016) and the significant alignments (false discovery rate < 0.05) were selected. The colors represent the nature of the miRNA-related locus (existing miRNA gene, protein-coding gene, *i.e* CDS and non-CDS, and transposable element). Non-CDS annotations include introns and untranslated regions. Bases belonging to more than one category (origins overlapping CDS and non-CDS in protein-coding genes) were added to each category.

**Table S1: Sources of origin of the 310 *A. halleri*-specific miRNA genes.** Each miRNA locus is described in a row, and defined by its genomic position. Each one is given a unique ID containing information about the family to which it belongs (letters indicate individual members of the family). Preferential loading into AGO proteins is detailed in Pavan et al. (2024) and is based on abundance in either of the two fractions relative to the input . “not loaded” indicates that the miRNA was observed in the input but neither in the AGO1 nor in the AGO4 fractions. “NA” indicates that the miRNA was not observed in the input. Hits to miRNA, TE and protein-coding genes or “unannotated sequences” are indicated, and the origin estimated for each of them is indicated in the last column.

| Loci | miRNA | Argonaute loading | Related miRNA | Related transposon superfamily | Related gene ID | Related unnanotated loci | Origin |
|---|---|---|---|---|---|---|---|
| ddAraHall_1.3_1:7097-7324 | aha-miR1a | AGO4 | . | Harbinger | . | . | TE |
| ddAraHall_1.3_1:597043-597210 | aha-miR2 | NA | . | . | . | . | no hit |
| ddAraHall_1.3_1:1110805-1110918 | aha-miR3 | AGO4 | . | . | . | ddAraHall_1.3_2:455744-455856 | unannotated |
| ddAraHall_1.3_1:4182510-4182588 | aha-miR4 | not loaded | . | . | . | . | no hit |
| ddAraHall_1.3_1:5970293-5970366 | aha-miR5 | AGO4 | . | . | . | . | no hit |
| ddAraHall_1.3_1:6550675-6551066 | aha-miR6 | NA | . | MuDR | . | . | TE |
| ddAraHall_1.3_1:6796987-6797284 | aha-miR7 | AGO4 | . | MITE | . | . | TE |
| ddAraHall_1.3_1:6926780-6926909 | aha-miR8a | not loaded | aha-miR8b | MuDR | Ah6g755439 | . | TE |
| ddAraHall_1.3_1:7150694-7150861 | aha-miR9 | not loaded | . | . | . | ddAraHall_1.3_4:16051370-16051537 | unannotated |
| ddAraHall_1.3_1:7994029-7994165 | aha-miR10a | AGO4 | aha-miR69b | MITE | . | . | TE |
| ddAraHall_1.3_1:9114798-9115187 | aha-miR11 | NA | . | . | . | . | no hit |
| ddAraHall_1.3_1:9764331-9764433 | aha-miR12 | AGO1 | . | . | . | . | no hit |
| ddAraHall_1.3_1:12476759-12476945 | aha-miR13a | AGO1 | . | MuDR | . | . | TE |
| ddAraHall_1.3_1:12478798-12478950 | aha-miR14 | NA | . | . | . | . | no hit |
| ddAraHall_1.3_1:12617460-12617856 | aha-miR15 | NA | . | . | . | ddAraHall_1.3_1:12617460-12617855 | unannotated |
| ddAraHall_1.3_1:12695938-12696235 | aha-miR16a | AGO1 | aha-miR16b | . | . | . | mirna |
| ddAraHall_1.3_1:12944744-12945222 | aha-miR17a | AGO4 | . | . | . | ddAraHall_1.3_1:5060401-5060683 | unannotated |
| ddAraHall_1.3_1:12970700-12970966 | aha-miR18 | AGO4 | . | . | . | ddAraHall_1.3_1:12048465-12048729 | unannotated |
| ddAraHall_1.3_1:13884475-13884654 | aha-miR19 | not loaded | . | . | . | ddAraHall_1.3_1:13878443-13878595 | unannotated |
| ddAraHall_1.3_1:14212519-14212831 | aha-miR20a | AGO4 | . | Helitron | . | . | TE |
| ddAraHall_1.3_1:15130841-15130959 | aha-miR17b | AGO4 | aha-miR17d | . | . | . | mirna |
| ddAraHall_1.3_1:15958417-15958786 | aha-miR21 | AGO4 | . | . | . | . | no hit |
| ddAraHall_1.3_1:16043382-16043504 | aha-miR22 | NA | . | . | . | . | no hit |
| ddAraHall_1.3_1:16108090-16108201 | aha-miR23 | AGO4 | . | . | . | ddAraHall_1.3_7:14550089-14550196 | unannotated |
| ddAraHall_1.3_1:17213304-17213403 | aha-miR24 | AGO1 | . | . | . | . | no hit |
| ddAraHall_1.3_1:17280778-17280979 | aha-miR25 | NA | . | . | . | ddAraHall_1.3_6:23790298-23790412 | unannotated |
| ddAraHall_1.3_1:18940212-18940293 | aha-miR26a | AGO4 | . | . | . | ddAraHall_1.3_1:26239175-26239253 | unannotated |
| ddAraHall_1.3_1:18949129-18949201 | aha-miR26b | AGO4 | aha-miR22c | . | . | . | mirna |
| ddAraHall_1.3_1:18965051-18965123 | aha-miR22c | AGO4 | aha-miR26b | . | . | . | mirna |
| ddAraHall_1.3_1:21082592-21082878 | aha-miR27a | NA | aha-miR27b | MITE | . | . | TE |
| ddAraHall_1.3_1:24193763-24193839 | aha-miR28 | not loaded | . | . | . | ddAraHall_1.3_3:2091472-2091547 | unannotated |
| ddAraHall_1.3_1:25661705-25661859 | aha-miR29 | AGO4 | . | . | . | . | no hit |
| ddAraHall_1.3_1:26496842-26496995 | aha-miR30 | AGO1 | . | . | . | . | no hit |
| ddAraHall_1.3_1:26743281-26743351 | aha-miR31 | NA | . | Helitron | . | . | TE |
| ddAraHall_1.3_1:27770076-27770170 | aha-miR32 | NA | . | . | . | . | no hit |
| ddAraHall_1.3_1:28527238-28527331 | aha-miR33 | AGO4 | . | MuDR | . | . | TE |
| ddAraHall_1.3_1:29121439-29121514 | aha-miR34 | NA | . | . | . | ddAraHall_1.3_1:11781723-11781797 | unannotated |
| ddAraHall_1.3_1:30231157-30231299 | aha-miR35 | AGO4 | . | . | . | ddAraHall_1.3_1:5652210-5652351 | unannotated |
| ddAraHall_1.3_1:30923864-30924036 | aha-miR36 | not loaded | . | MITE | Ah7g820024 | . | TE |
| ddAraHall_1.3_2:654836-654972 | aha-miR10b | AGO4 | aha-miR10c | hAT | . | . | TE |
| ddAraHall_1.3_2:1040654-1040838 | aha-miR37 | AGO1 | . | . | . | . | no hit |
| ddAraHall_1.3_2:1079998-1080127 | aha-miR38 | NA | . | . | . | ddAraHall_1.3_2:1080011-1080114 | unannotated |
| ddAraHall_1.3_2:1165514-1165830 | aha-miR39 | NA | . | Helitron | Ah2g788505 | . | TE |
| ddAraHall_1.3_2:1353780-1353919 | aha-miR40 | AGO1 | . | . | . | . | no hit |
| ddAraHall_1.3_2:1400850-1401248 | aha-miR41 | AGO4 | . | MITE | . | . | TE |
| ddAraHall_1.3_2:1657040-1657266 | aha-miR42 | AGO1 | . | Helitron | Ah2g788816 | . | TE |
| ddAraHall_1.3_2:2407586-2407833 | aha-miR43 | NA | . | . | . | . | no hit |
| ddAraHall_1.3_2:2574865-2575068 | aha-miR44 | NA | . | . | Ah8g677304 | . | CDS |
| ddAraHall_1.3_2:2941148-2941306 | aha-miR45 | NA | . | Copia | . | . | TE |
| ddAraHall_1.3_2:3242554-3242810 | aha-miR46 | NA | . | . | . | . | no hit |
| ddAraHall_1.3_2:3829873-3829970 | aha-miR47 | NA | . | . | . | . | no hit |
| ddAraHall_1.3_2:4234679-4234933 | aha-miR48 | NA | . | MuDR | . | . | TE |
| ddAraHall_1.3_2:4540022-4540180 | aha-miR49 | AGO4 | . | Helitron | . | . | TE |
| ddAraHall_1.3_2:7018253-7018341 | aha-miR50a | not loaded | aha-miR50b | Tase | . | . | TE |
| ddAraHall_1.3_2:8104715-8104794 | aha-miR51 | NA | . | . | . | . | no hit |
| ddAraHall_1.3_2:8489772-8489964 | aha-miR13b | AGO1 | . | MuDR | . | . | TE |
| ddAraHall_1.3_2:8724691-8725066 | aha-miR52b | AGO4 | aha-miR52a | hAT | . | . | TE |
| ddAraHall_1.3_2:8882974-8883276 | aha-miR52c | AGO4 | aha-miR52c | hAT | . | . | TE |
| ddAraHall_1.3_2:12282574-12282660 | aha-miR53a | AGO1 AGO4 | aha-miR53b | . | . | . | mirna |
| ddAraHall_1.3_2:12297552-12297638 | aha-miR54 | AGO1 | . | . | Ah2g796705 | . | non_CDS |
| ddAraHall_1.3_2:12305160-12305231 | aha-miR55 | not loaded | . | . | Ah2g796659 | . | CDS |
| ddAraHall_1.3_2:12387224-12387308 | aha-miR53b | AGO1 AGO4 | aha-miR53a | . | . | . | mirna |
| ddAraHall_1.3_2:12442239-12442523 | aha-miR56 | AGO4 | . | . | . | . | no hit |
| ddAraHall_1.3_2:12521689-12521770 | aha-miR57 | not loaded | . | Gypsy | . | . | TE |
| ddAraHall_1.3_2:12728661-12728791 | aha-miR58 | NA | . | . | Ah7g817482 | . | CDS |
| ddAraHall_1.3_2:13979848-13980108 | aha-miR59 | NA | . | . | Ah2g797910 | . | non_CDS |
| ddAraHall_1.3_2:14131833-14132128 | aha-miR60a | NA | . | classII | Ah4g698843 | . | TE |
| ddAraHall_1.3_2:14643263-14643372 | aha-miR61 | AGO1 | . | . | . | ddAraHall_1.3_57:53109-53216 | unannotated |
| ddAraHall_1.3_2:14761433-14761596 | aha-miR62 | not loaded | . | MuDR | . | . | TE |
| ddAraHall_1.3_2:16239944-16240336 | aha-miR63 | not loaded | . | MITE | . | . | TE |
| ddAraHall_1.3_2:16897508-16897631 | aha-miR64 | AGO1 | . | . | . | . | no hit |
| ddAraHall_1.3_2:17223468-17223620 | aha-miR65 | AGO4 | . | . | . | ddAraHall_1.3_4:3971309-3971401 | unannotated |
| ddAraHall_1.3_2:17356391-17356790 | aha-miR66 | NA | . | Harbinger | . | . | TE |
| ddAraHall_1.3_2:17773794-17773864 | aha-miR67 | NA | . | LINE | . | . | TE |
| ddAraHall_1.3_2:17819501-17819577 | aha-miR68 | NA | . | . | . | . | no hit |
| ddAraHall_1.3_2:18985108-18985352 | aha-miR69a | not loaded | aha-miR10b | . | . | . | mirna |
| ddAraHall_1.3_2:19336888-19337286 | aha-miR70 | NA | . | MITE | . | . | TE |
| ddAraHall_1.3_2:19769962-19770098 | aha-miR10c | AGO4 | aha-miR10b | MITE | . | . | TE |
| ddAraHall_1.3_2:20183160-20183232 | aha-miR71 | AGO4 | . | SINE | . | . | TE |
| ddAraHall_1.3_2:20577686-20578077 | aha-miR72 | AGO1 | . | . | Ah2g806442 | . | non_CDS |
| ddAraHall_1.3_2:21045619-21045873 | aha-miR73 | AGO1 | . | LINE | . | . | TE |
| ddAraHall_1.3_2:21366920-21367085 | aha-miR74 | AGO4 | . | . | . | ddAraHall_1.3_6:2600688-2600861 | unannotated |
| ddAraHall_1.3_2:22168204-22168422 | aha-miR75 | NA | . | . | . | . | no hit |
| ddAraHall_1.3_2:22274527-22274687 | aha-miR76 | AGO4 | . | . | . | ddAraHall_1.3_8:791106-791178 | unannotated |
| ddAraHall_1.3_2:22344000-22344399 | aha-miR77a | AGO4 | . | MuDR | . | . | TE |
| ddAraHall_1.3_2:22914344-22914527 | aha-miR78 | AGO1 | . | . | . | . | no hit |
| ddAraHall_1.3_2:24134806-24134876 | aha-miR79 | NA | . | LINE | . | . | TE |
| ddAraHall_1.3_3:50804-50900 | aha-miR80 | not loaded | . | . | . | . | no hit |
| ddAraHall_1.3_3:380856-381100 | aha-miR81 | AGO1 and AGO4 | . | . | Ah3g756269 | . | non_CDS |
| ddAraHall_1.3_3:489526-489599 | aha-miR82 | AGO4 | . | . | . | . | no hit |
| ddAraHall_1.3_3:1181997-1182232 | aha-miR83 | AGO1 | . | . | . | . | no hit |
| ddAraHall_1.3_3:1519720-1520111 | aha-miR84a | AGO1 | . | MuDR | . | . | TE |
| ddAraHall_1.3_3:2522442-2522528 | aha-miR85 | NA | . | MITE | . | . | TE |
| ddAraHall_1.3_3:3245548-3245945 | aha-miR86 | AGO4 | . | . | . | . | no hit |
| ddAraHall_1.3_3:4038590-4038701 | aha-miR87 | AGO4 | . | . | . | ddAraHall_1.3_2:7563621-7563731 | unannotated |
| ddAraHall_1.3_3:4742349-4742429 | aha-miR88 | AGO4 | . | Mariner | . | . | TE |
| ddAraHall_1.3_3:5327411-5327644 | aha-miR89a | AGO4 | . | MITE | . | . | TE |
| ddAraHall_1.3_3:6876400-6876756 | aha-miR90 | AGO1 | . | . | Ah3g769079 | . | non_CDS |
| ddAraHall_1.3_3:7131758-7131914 | aha-miR91 | AGO1 | . | . | . | . | no hit |

| | | | | | | | |
|---|---|---|---|---|---|---|---|
| ddAraHall_1.3_3:7285651-7285798 | aha-miR92 | NA | . | . | . | . | no hit |
| ddAraHall_1.3_3:7289322-7289456 | aha-miR93 | NA | . | . | . | . | no hit |
| ddAraHall_1.3_3:8354471-8354564 | aha-miR94a | NA | aha-miR94b | Helitron | . | . | TE |
| ddAraHall_1.3_3:9066414-9066813 | aha-miR95 | AGO1 | . | . | Ah3g772753 | . | CDS and non_CDS |
| ddAraHall_1.3_3:10162772-10162901 | aha-miR96 | AGO4 | . | MuDR | . | . | TE |
| ddAraHall_1.3_3:10898566-10898829 | aha-miR97 | NA | . | . | . | ddAraHall_1.3_3:11999888-12000150 | unannotated |
| ddAraHall_1.3_3:13770887-13770961 | aha-miR98 | AGO4 | . | . | Ah6g753650 | . | non_CDS |
| ddAraHall_1.3_3:14399138-14399208 | aha-miR99 | NA | . | . | Ah1g859091 | . | CDS |
| ddAraHall_1.3_3:14523035-14523106 | aha-miR100 | NA | . | Harbinger | . | . | TE |
| ddAraHall_1.3_3:14877193-14877356 | aha-miR13c | NA | . | MuDR | . | . | TE |
| ddAraHall_1.3_3:15036899-15037002 | aha-miR101 | AGO4 | . | . | . | ddAraHall_1.3_2:4733820-4733917 | unannotated |
| ddAraHall_1.3_3:16074864-16074958 | aha-miR102 | AGO4 | . | . | Ah8g681064 | . | non_CDS |
| ddAraHall_1.3_3:21006817-21006966 | aha-miR103 | AGO4 | . | MITE | . | . | TE |
| ddAraHall_1.3_3:21433207-21433591 | aha-miR104 | AGO4 | . | TRIM_LARD | . | . | TE |
| ddAraHall_1.3_3:24698510-24698595 | aha-miR105 | AGO4 | . | . | . | . | no hit |
| ddAraHall_1.3_3:26426776-26426927 | aha-miR106 | AGO1 | . | . | . | . | no hit |
| ddAraHall_1.3_4:2533-2620 | aha-miR107 | AGO4 | . | . | . | ddAraHall_1.3_113:34708-34793 | unannotated |
| ddAraHall_1.3_4:16548-16634 | aha-miR108 | AGO1 | . | . | . | ddAraHall_1.3_113:28886-28971 | unannotated |
| ddAraHall_1.3_4:1692116-1692201 | aha-miR109 | AGO4 | . | MuDR | . | . | TE |
| ddAraHall_1.3_4:2193128-2193206 | aha-miR110 | NA | . | MuDR | . | . | TE |
| ddAraHall_1.3_4:2237258-2237333 | aha-miR111 | not loaded | . | . | . | ddAraHall_1.3_2:23394454-23394528 | unannotated |
| ddAraHall_1.3_4:2694679-2694776 | aha-miR112 | NA | . | hAT | . | . | TE |
| ddAraHall_1.3_4:2951034-2951433 | aha-miR113 | AGO1 | . | Gypsy | . | . | TE |
| ddAraHall_1.3_4:3115375-3115469 | aha-miR114 | AGO1 | . | MuDR | . | . | TE |
| ddAraHall_1.3_4:3363683-3364082 | aha-miR115 | AGO4 | . | . | . | ddAraHall_1.3_8:3349760-3350050 | unannotated |
| ddAraHall_1.3_4:3467940-3468052 | aha-miR116 | AGO1 | . | . | . | . | no hit |
| ddAraHall_1.3_4:3584586-3584733 | aha-miR117a | AGO4 | . | MITE | . | . | TE |
| ddAraHall_1.3_4:3873421-3873551 | aha-miR13d | AGO4 | . | MuDR | . | . | TE |
| ddAraHall_1.3_4:4223984-4224073 | aha-miR118 | NA | . | . | . | . | no hit |
| ddAraHall_1.3_4:4969248-4969316 | aha-miR119 | NA | . | classII | . | . | TE |
| ddAraHall_1.3_4:6772792-6772950 | aha-miR120 | NA | . | Helitron | . | . | TE |
| ddAraHall_1.3_4:12455573-12455769 | aha-miR121 | AGO1 | . | Gypsy | . | . | TE |
| ddAraHall_1.3_4:12505209-12505461 | aha-miR122 | AGO1 | . | . | . | . | no hit |
| ddAraHall_1.3_4:15489381-15489638 | aha-miR16b | AGO1 and AGO4 | aha-miR16a | . | Ah1g871602 | . | non_CDS |
| ddAraHall_1.3_4:17255266-17255628 | aha-miR123 | NA | . | MuDR | . | . | TE |
| ddAraHall_1.3_4:17391815-17391955 | aha-miR124 | not loaded | . | classII | . | . | TE |
| ddAraHall_1.3_4:18812318-18812402 | aha-miR125 | NA | . | . | . | . | no hit |
| ddAraHall_1.3_4:19314274-19314366 | aha-miR126a | AGO4 | . | Helitron | . | . | TE |
| ddAraHall_1.3_4:19820795-19821194 | aha-miR127 | AGO1 | . | . | . | ddAraHall_1.3_4:19820795-19821191 | unannotated |
| ddAraHall_1.3_4:19972603-19973002 | aha-miR128 | AGO4 | . | . | . | . | no hit |
| ddAraHall_1.3_4:20889608-20889681 | aha-miR129 | AGO4 | . | . | . | ddAraHall_1.3_4:12407325-12407397 | unannotated |
| ddAraHall_1.3_4:21169354-21169457 | aha-miR130 | NA | . | . | . | . | no hit |
| ddAraHall_1.3_4:21249542-21249798 | aha-miR131 | NA | . | . | . | . | no hit |
| ddAraHall_1.3_4:21628295-21628686 | aha-miR84b | NA | . | MuDR | . | . | TE |
| ddAraHall_1.3_4:22217543-22217924 | aha-miR1b | AGO4 | aha-miR1a | Harbinger | . | . | TE |
| ddAraHall_1.3_4:25587214-25587350 | aha-miR132 | not loaded | . | . | . | ddAraHall_1.3_2:2879260-2879383 | unannotated |
| ddAraHall_1.3_4:25672130-25672298 | aha-miR133 | NA | . | . | . | . | no hit |
| ddAraHall_1.3_5:799702-799812 | aha-miR134 | AGO1 | . | . | . | . | no hit |
| ddAraHall_1.3_5:4033392-4033464 | aha-miR135 | AGO4 | . | . | . | . | no hit |
| ddAraHall_1.3_5:4082060-4082306 | aha-miR136 | AGO4 | . | LINE | . | . | TE |
| ddAraHall_1.3_5:4743481-4743756 | aha-miR17c | AGO4 | . | . | . | ddAraHall_1.3_1:5060403-5060677 | unannotated |
| ddAraHall_1.3_5:5100924-5101225 | aha-miR137 | NA | . | MuDR | . | . | TE |
| ddAraHall_1.3_5:5187768-5187868 | aha-miR138 | AGO1 | . | . | . | . | no hit |
| ddAraHall_1.3_5:5595979-5596075 | aha-miR139a | AGO4 | . | Harbinger | . | . | TE |
| ddAraHall_1.3_5:6706449-6706540 | aha-miR140 | AGO4 | . | Chimeric | . | . | TE |
| ddAraHall_1.3_5:6888153-6888471 | aha-miR141 | AGO1 and AGO4 | . | . | . | ddAraHall_1.3_4:2276068-2276370 | unannotated |
| ddAraHall_1.3_5:7407494-7407615 | aha-miR142 | NA | . | . | . | . | no hit |
| ddAraHall_1.3_5:7503658-7504019 | aha-miR143a | AGO1 | aha-miR143b | . | . | . | mirna |
| ddAraHall_1.3_5:7728499-7728592 | aha-miR144 | AGO4 | . | . | . | ddAraHall_1.3_6:13320871-13320959 | unannotated |
| ddAraHall_1.3_5:8031750-8031935 | aha-miR145 | AGO1 | . | . | . | . | no hit |
| ddAraHall_1.3_5:8032006-8032190 | aha-miR146 | AGO1 and AGO4 | . | . | . | . | no hit |
| ddAraHall_1.3_5:8782553-8782730 | aha-miR147 | AGO1 | . | . | . | . | no hit |
| ddAraHall_1.3_5:8867567-8867760 | aha-miR148 | AGO4 | . | Mariner | . | . | TE |
| ddAraHall_1.3_5:9431151-9431336 | aha-miR149 | AGO1 | . | . | . | ddAraHall_1.3_5:9431160-9431328 | unannotated |
| ddAraHall_1.3_5:10212744-10212846 | aha-miR150 | not loaded | . | . | Ah2g796238 | . | non_CDS |
| ddAraHall_1.3_5:11286743-11287141 | aha-miR77b | AGO1 and AGO4 | . | MuDR | . | . | TE |
| ddAraHall_1.3_5:11765922-11766032 | aha-miR151 | AGO4 | . | Copia | . | . | TE |
| ddAraHall_1.3_5:14836566-14836635 | aha-miR152 | AGO4 | . | Gypsy | . | . | TE |
| ddAraHall_1.3_5:15128633-15128961 | aha-miR153 | NA | . | hAT | . | . | TE |
| ddAraHall_1.3_5:16669714-16670057 | aha-miR154 | AGO4 | . | Harbinger | Ah3g780861 | . | TE |
| ddAraHall_1.3_5:16670340-16670411 | aha-miR155 | NA | . | Harbinger | Ah3g781278 | . | TE |
| ddAraHall_1.3_5:17098697-17098767 | aha-miR156a | AGO4 | . | Gypsy | . | . | TE |
| ddAraHall_1.3_5:17104660-17104730 | aha-miR156b | AGO4 | . | Gypsy | . | . | TE |
| ddAraHall_1.3_5:17548540-17548912 | aha-miR13e | NA | . | MuDR | . | . | TE |
| ddAraHall_1.3_5:17658696-17658794 | aha-miR157a | AGO4 | aha-miR157c | . | . | . | mirna |
| ddAraHall_1.3_5:17696230-17696328 | aha-miR157b | AGO4 | aha-miR157c | . | . | . | mirna |
| ddAraHall_1.3_5:17731821-17731912 | aha-miR158 | NA | . | MITE | . | . | TE |
| ddAraHall_1.3_5:18377809-18378006 | aha-miR159 | AGO4 | . | CACTA | Ah2g787547 | . | TE |
| ddAraHall_1.3_5:18668527-18668787 | aha-miR160 | AGO1 | . | Copia | . | . | TE |
| ddAraHall_1.3_5:18743482-18743610 | aha-miR161 | NA | . | . | . | ddAraHall_1.3_8:11273053-11273126 | unannotated |
| ddAraHall_1.3_5:19794624-19794853 | aha-miR162 | AGO1 | . | . | . | . | no hit |
| ddAraHall_1.3_5:20201081-20201341 | aha-miR163 | NA | . | . | . | ddAraHall_1.3_4:16051399-16051509 | unannotated |
| ddAraHall_1.3_5:21454319-21454411 | aha-miR164 | not loaded | . | classII | . | . | TE |
| ddAraHall_1.3_5:22173663-22173782 | aha-miR165 | not loaded | . | . | . | . | no hit |
| ddAraHall_1.3_6:212049-212147 | aha-miR157c | AGO4 | aha-miR157a | . | . | . | mirna |
| ddAraHall_1.3_6:1183017-1183416 | aha-miR166a | AGO1 | . | MuDR | . | . | TE |
| ddAraHall_1.3_6:1429141-1429476 | aha-miR167 | AGO1 | . | . | . | ddAraHall_1.3_6:1429141-1429470 | unannotated |
| ddAraHall_1.3_6:1581376-1581679 | aha-miR168 | AGO4 | . | . | Ah6g726079 | . | CDS and non_CDS |
| ddAraHall_1.3_6:1824783-1824862 | aha-miR169 | NA | . | . | . | . | no hit |
| ddAraHall_1.3_6:3717935-3718007 | aha-miR139b | AGO4 | aha-miR139a | Harbinger | . | . | TE |
| ddAraHall_1.3_6:3856282-3856674 | aha-miR170 | AGO4 | . | . | . | ddAraHall_1.3_6:3856282-3856673 | unannotated |
| ddAraHall_1.3_6:4988724-4988808 | aha-miR171 | not loaded | . | . | . | . | no hit |
| ddAraHall_1.3_6:5089955-5090080 | aha-miR172 | NA | . | . | . | . | no hit |
| ddAraHall_1.3_6:5692368-5692661 | aha-miR173 | AGO4 | . | . | Ah2g805320 | . | non_CDS |
| ddAraHall_1.3_6:5935179-5935557 | aha-miR174 | NA | . | . | . | . | no hit |
| ddAraHall_1.3_6:6879442-6879630 | aha-miR175 | AGO1 | . | . | . | . | no hit |
| ddAraHall_1.3_6:7540165-7540337 | aha-miR176 | NA | . | . | . | . | no hit |
| ddAraHall_1.3_6:10789907-10790097 | aha-miR177 | AGO4 | . | MuDR | . | . | TE |
| ddAraHall_1.3_6:11072188-11072587 | aha-miR178 | AGO4 | . | MuDR | . | . | TE |
| ddAraHall_1.3_6:11183398-11183475 | aha-miR179 | AGO4 | . | . | . | ddAraHall_1.3_1:12609199-12609267 | unannotated |

| | | | | | | | |
|---|---|---|---|---|---|---|---|
| ddAraHall_1.3_6:11517954-11518353 | aha-miR166b | AGO1 | . | MuDR | . | . | TE |
| ddAraHall_1.3_6:12139312-12139463 | aha-miR180 | NA | . | CACTA | . | . | TE |
| ddAraHall_1.3_6:13275826-13275899 | aha-miR181 | not loaded | . | LINE | . | . | TE |
| ddAraHall_1.3_6:13609057-13609365 | aha-miR143b | AGO1 | . | . | . | ddAraHall_1.3_5:21848306-21848612 | unannotated |
| ddAraHall_1.3_6:16512027-16512115 | aha-miR50b | not loaded | aha-miR50a | Tase | . | . | TE |
| ddAraHall_1.3_6:19777638-19777707 | aha-miR182a | AGO4 | . | Harbinger | . | . | TE |
| ddAraHall_1.3_6:19932709-19932843 | aha-miR183 | not loaded | . | . | . | ddAraHall_1.3_2:7207516-7207648 | unannotated |
| ddAraHall_1.3_6:20058647-20058741 | aha-miR184 | NA | . | Gypsy | . | . | TE |
| ddAraHall_1.3_6:20635169-20635565 | aha-miR185 | AGO1 | . | Gypsy | . | . | TE |
| ddAraHall_1.3_6:20636173-20636298 | aha-miR186 | NA | . | Gypsy | . | . | TE |
| ddAraHall_1.3_6:21108871-21109269 | aha-miR187 | AGO1 | . | . | Ah6g748522 | . | non_CDS |
| ddAraHall_1.3_6:22098938-22099051 | aha-miR188 | NA | . | LINE | . | . | TE |
| ddAraHall_1.3_6:22530527-22530604 | aha-miR189 | NA | . | . | . | . | no hit |
| ddAraHall_1.3_6:23366990-23367141 | aha-miR190 | NA | . | . | . | . | no hit |
| ddAraHall_1.3_6:23493380-23493500 | aha-miR13f | AGO4 | aha-miR13d | . | . | . | mirna |
| ddAraHall_1.3_6:23718664-23718807 | aha-miR191 | AGO1 | . | . | . | ddAraHall_1.3_6:23719580-23719724 | unannotated |
| ddAraHall_1.3_6:23718994-23719393 | aha-miR192 | AGO4 | . | . | . | ddAraHall_1.3_6:23718994-23719392 | unannotated |
| ddAraHall_1.3_6:24140858-24141007 | aha-miR193 | NA | . | . | . | ddAraHall_1.3_1:25337996-25338105 | unannotated |
| ddAraHall_1.3_6:25358692-25358872 | aha-miR194 | NA | . | . | . | . | no hit |
| ddAraHall_1.3_6:25850330-25850401 | aha-miR195 | NA | . | . | . | . | no hit |
| ddAraHall_1.3_6:25899878-25900066 | aha-miR196 | AGO4 | . | . | . | ddAraHall_1.3_7:13346758-13346869 | unannotated |
| ddAraHall_1.3_6:26579278-26579412 | aha-miR8b | not loaded | aha-miR8a | MuDR | . | . | TE |
| ddAraHall_1.3_7:696100-696320 | aha-miR197 | NA | . | . | . | . | no hit |
| ddAraHall_1.3_7:1128578-1128666 | aha-miR198a | not loaded | aha-miR198b | MITE | Ah7g814814 | . | TE |
| ddAraHall_1.3_7:1134534-1134622 | aha-miR198b | not loaded | aha-miR198a | MITE | Ah7g814815 | . | TE |
| ddAraHall_1.3_7:1393035-1393252 | aha-miR199 | NA | . | MuDR | . | . | TE |
| ddAraHall_1.3_7:2629705-2629774 | aha-miR200 | AGO4 | . | hAT | Ah2g787664 | . | TE |
| ddAraHall_1.3_7:3288753-3289152 | aha-miR201 | AGO1 | . | Gypsy | Ah1g881161 | . | TE |
| ddAraHall_1.3_7:3576864-3577040 | aha-miR28b | AGO4 | . | Gypsy | . | . | TE |
| ddAraHall_1.3_7:3643881-3644280 | aha-miR202 | NA | . | . | . | ddAraHall_1.3_4:6001719-6002065 | unannotated |
| ddAraHall_1.3_7:3930157-3930325 | aha-miR203 | not loaded | . | . | . | . | no hit |
| ddAraHall_1.3_7:4445689-4446037 | aha-miR204 | NA | . | . | . | ddAraHall_1.3_7:4447728-4448075 | unannotated |
| ddAraHall_1.3_7:4915341-4915740 | aha-miR205 | AGO1 | . | . | . | . | no hit |
| ddAraHall_1.3_7:4922175-4922434 | aha-miR206 | NA | . | . | . | ddAraHall_1.3_6:15446813-15447070 | unannotated |
| ddAraHall_1.3_7:5132326-5132441 | aha-miR207 | NA | . | . | . | ddAraHall_1.3_7:5141227-5141341 | unannotated |
| ddAraHall_1.3_7:5351752-5352040 | aha-miR27b | NA | aha-miR27a | MITE | . | . | TE |
| ddAraHall_1.3_7:5657426-5657559 | aha-miR208 | NA | . | . | . | ddAraHall_1.3_4:25637450-25637584 | unannotated |
| ddAraHall_1.3_7:5781414-5781561 | aha-miR209 | AGO1 | . | Copia | . | . | TE |
| ddAraHall_1.3_7:5787104-5787503 | aha-miR210 | AGO1 | . | Gypsy | . | . | TE |
| ddAraHall_1.3_7:6468310-6468415 | aha-miR211 | NA | . | . | . | . | no hit |
| ddAraHall_1.3_7:8846455-8846651 | aha-miR212a | AGO1 | aha-miR212d | . | Ah7g840513 | . | non_CDS |
| ddAraHall_1.3_7:8917234-8917632 | aha-miR213 | AGO4 | . | Copia | Ah8g681966 | . | TE |
| ddAraHall_1.3_7:10187186-10187277 | aha-miR214 | AGO1 | . | CACTA | . | . | TE |
| ddAraHall_1.3_7:10346384-10346463 | aha-miR215 | AGO4 | . | Helitron | . | . | TE |
| ddAraHall_1.3_7:10362118-10362514 | aha-miR216a | NA | . | . | . | ddAraHall_1.3_7:10287187-10287293 | unannotated |
| ddAraHall_1.3_7:10446838-10447234 | aha-miR216b | NA | . | . | . | ddAraHall_1.3_7:10287187-10287293 | unannotated |
| ddAraHall_1.3_7:10468211-10468609 | aha-miR217 | NA | . | . | Ah7g813868 | . | CDS |
| ddAraHall_1.3_7:10492671-10493061 | aha-miR218 | NA | . | . | . | . | no hit |
| ddAraHall_1.3_7:10547665-10547917 | aha-miR219 | NA | . | . | . | . | no hit |
| ddAraHall_1.3_7:10746955-10747323 | aha-miR220 | AGO4 | . | Mariner | . | . | TE |
| ddAraHall_1.3_7:11066865-11067259 | aha-miR221 | NA | . | Harbinger | . | . | TE |
| ddAraHall_1.3_7:11583047-11583142 | aha-miR126b | AGO4 | . | Helitron | . | . | TE |
| ddAraHall_1.3_7:11584825-11584945 | aha-miR222 | AGO1 | . | . | . | ddAraHall_1.3_5:15019154-15019270 | unannotated |
| ddAraHall_1.3_7:12856721-12856970 | aha-miR223a | AGO4 | . | LINE | . | . | TE |
| ddAraHall_1.3_7:12880993-12881071 | aha-miR224 | NA | . | . | . | . | no hit |
| ddAraHall_1.3_7:13377081-13377370 | aha-miR225 | not loaded | . | . | . | ddAraHall_1.3_3:13724454-13724733 | unannotated |
| ddAraHall_1.3_7:13680752-13680872 | aha-miR13g | AGO4 | . | MuDR | . | . | TE |
| ddAraHall_1.3_7:14409493-14409710 | aha-miR226 | AGO4 | . | Mariner | . | . | TE |
| ddAraHall_1.3_7:15590528-15590679 | aha-miR227 | NA | . | . | . | ddAraHall_1.3_7:20527775-20527925 | unannotated |
| ddAraHall_1.3_7:15969166-15969565 | aha-miR228 | NA | . | . | . | ddAraHall_1.3_7:15969166-15969564 | unannotated |
| ddAraHall_1.3_7:16276716-16276897 | aha-miR229 | AGO1 | . | . | . | . | no hit |
| ddAraHall_1.3_7:16888813-16888989 | aha-miR230 | AGO1 | . | . | . | . | no hit |
| ddAraHall_1.3_7:17264581-17264667 | aha-miR231 | NA | . | . | . | . | no hit |
| ddAraHall_1.3_7:17898939-17899325 | aha-miR232 | AGO4 | . | MITE | . | . | TE |
| ddAraHall_1.3_7:19048671-19048831 | aha-miR233 | AGO4 | aha-miR117a | MITE | . | . | TE |
| ddAraHall_1.3_7:20352154-20352230 | aha-miR234 | AGO4 | . | . | . | ddAraHall_1.3_7:20351281-20351355 | unannotated |
| ddAraHall_1.3_7:20646337-20646443 | aha-miR235 | AGO1 | . | . | . | . | no hit |
| ddAraHall_1.3_7:21581567-21581647 | aha-miR236 | AGO4 | . | SINE | . | . | TE |
| ddAraHall_1.3_7:21594891-21595058 | aha-miR237 | AGO1 | . | . | . | . | no hit |
| ddAraHall_1.3_7:21661551-21661696 | aha-miR212b | AGO1 | aha-miR212c | . | . | . | mirna |
| ddAraHall_1.3_7:21684820-21684965 | aha-miR212c | AGO1 | aha-miR212b | . | Ah7g837326 | . | non_CDS |
| ddAraHall_1.3_7:23323053-23323272 | aha-miR212d | AGO1 | . | . | Ah7g840517 | . | non_CDS |
| ddAraHall_1.3_7:23491711-23492022 | aha-miR69b | not loaded | aha-miR69c | . | . | . | mirna |
| ddAraHall_1.3_7:25915414-25915560 | aha-miR238 | AGO1 | . | . | Ah7g844770 | . | non_CDS |
| ddAraHall_1.3_8:1308929-1309274 | aha-miR239 | not loaded | . | Helitron | . | . | TE |
| ddAraHall_1.3_8:2258793-2259095 | aha-miR223b | not loaded | . | LINE | . | . | TE |
| ddAraHall_1.3_8:2871678-2871800 | aha-miR182b | AGO4 | . | . | . | ddAraHall_1.3_6:11782402-11782523 | unannotated |
| ddAraHall_1.3_8:3254822-3254904 | aha-miR240 | NA | . | . | . | . | no hit |
| ddAraHall_1.3_8:3464701-3464771 | aha-miR241 | NA | . | MuDR | . | . | TE |
| ddAraHall_1.3_8:4569074-4569144 | aha-miR242 | not loaded | . | . | . | . | no hit |
| ddAraHall_1.3_8:4744148-4744414 | aha-miR69c | AGO4 | . | . | . | ddAraHall_1.3_3:13030525-13030785 | unannotated |
| ddAraHall_1.3_8:5414175-5414485 | aha-miR77c | AGO4 | . | MuDR | . | . | TE |
| ddAraHall_1.3_8:5469657-5469843 | aha-miR13h | AGO1 | . | MuDR | . | . | TE |
| ddAraHall_1.3_8:5804212-5804280 | aha-miR243 | NA | . | . | . | . | no hit |
| ddAraHall_1.3_8:6971012-6971119 | aha-miR244a | NA | aha-miR244b | Helitron | . | . | TE |
| ddAraHall_1.3_8:6975673-6975780 | aha-miR244b | NA | aha-miR244a | Helitron | . | . | TE |
| ddAraHall_1.3_8:7170316-7170715 | aha-miR245 | NA | . | . | Ah8g676827 | . | CDS |
| ddAraHall_1.3_8:8695615-8695796 | aha-miR246 | AGO1 | . | . | . | . | no hit |
| ddAraHall_1.3_8:9061231-9061630 | aha-miR247 | AGO1 | . | Harbinger | . | . | TE |
| ddAraHall_1.3_8:9496290-9496457 | aha-miR248 | AGO4 | . | MuDR | Ah7g813974 | . | TE |
| ddAraHall_1.3_8:9660035-9660128 | aha-miR94b | NA | aha-miR94a | Helitron | . | . | TE |
| ddAraHall_1.3_8:9669316-9669402 | aha-miR20b | AGO4 | . | . | . | ddAraHall_1.3_1:6126278-6126359 | unannotated |
| ddAraHall_1.3_8:11214846-11215102 | aha-miR60b | NA | . | classII | Ah4g698843 | . | TE |
| ddAraHall_1.3_8:11440574-11440688 | aha-miR249 | AGO4 | . | . | . | ddAraHall_1.3_7:22964376-22964490 | unannotated |
| ddAraHall_1.3_8:11810338-11810417 | aha-miR250 | AGO1 | . | MuDR | . | . | TE |
| ddAraHall_1.3_8:12771972-12772195 | aha-miR251 | NA | . | . | . | . | no hit |
| ddAraHall_1.3_8:13768769-13769168 | aha-miR252 | AGO4 | . | MITE | . | . | TE |
| ddAraHall_1.3_8:14169929-14170322 | aha-miR84c | NA | . | MuDR | . | . | TE |
| ddAraHall_1.3_8:16109880-16110253 | aha-miR253 | AGO4 | . | MITE | . | . | TE |
| ddAraHall_1.3_8:16420568-16420731 | aha-miR254 | AGO1 | . | . | . | . | no hit |

| | | | | | | | |
|---|---|---|---|---|---|---|---|
| ddAraHall_1.3_8:16644499-16644606 | aha-miR255 | AGO1 | . | . | . | ddAraHall_1.3_8:13352940-13352992 | unannotated |
| ddAraHall_1.3_8:17090518-17090803 | aha-miR256 | AGO1 | . | . | Ah8g686378 | . | non_CDS |
| ddAraHall_1.3_8:17090921-17091040 | aha-miR257 | NA | . | . | . | . | no hit |
| ddAraHall_1.3_8:17799486-17799817 | aha-miR258 | AGO1 | . | . | Ah8g687736 | . | non_CDS |
| ddAraHall_1.3_8:18031457-18031627 | aha-miR1c | NA | . | . | . | ddAraHall_1.3_6:11101719-11101885 | unannotated |
| ddAraHall_1.3_8:18133488-18133641 | aha-miR259 | AGO1 | . | . | . | . | no hit |
| ddAraHall_1.3_8:20021935-20022193 | aha-miR17d | not loaded | . | . | . | ddAraHall_1.3_7:19654918-19655175 | unannotated |
| ddAraHall_1.3_8:20669057-20669358 | aha-miR260 | NA | . | . | . | ddAraHall_1.3_57:88858-89008 | unannotated |
| ddAraHall_1.3_8:21283278-21283560 | aha-miR17e | NA | aha-miR17a | Harbinger | . | . | TE |
| ddAraHall_1.3_8:21793206-21793533 | aha-miR261 | NA | . | . | . | . | no hit |
| ddAraHall_1.3_8:22300087-22300233 | aha-miR262 | AGO4 | . | . | . | ddAraHall_1.3_6:5516899-5517046 | unannotated |

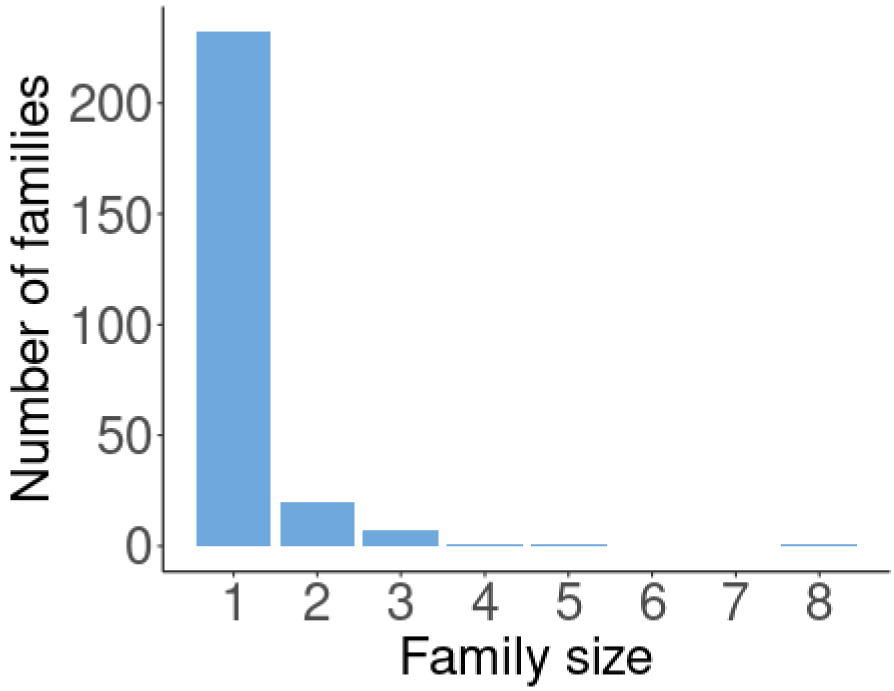


**Figure S2: Distribution of family sizes of *A. halleri*-specific miRNA genes**. Most miRNA genes belong to "singletons" (*i.e.* families with a single member). One family (Aha-miR13) comprises eight paralogs and likely derives from a MuDR TE (Figure 3).

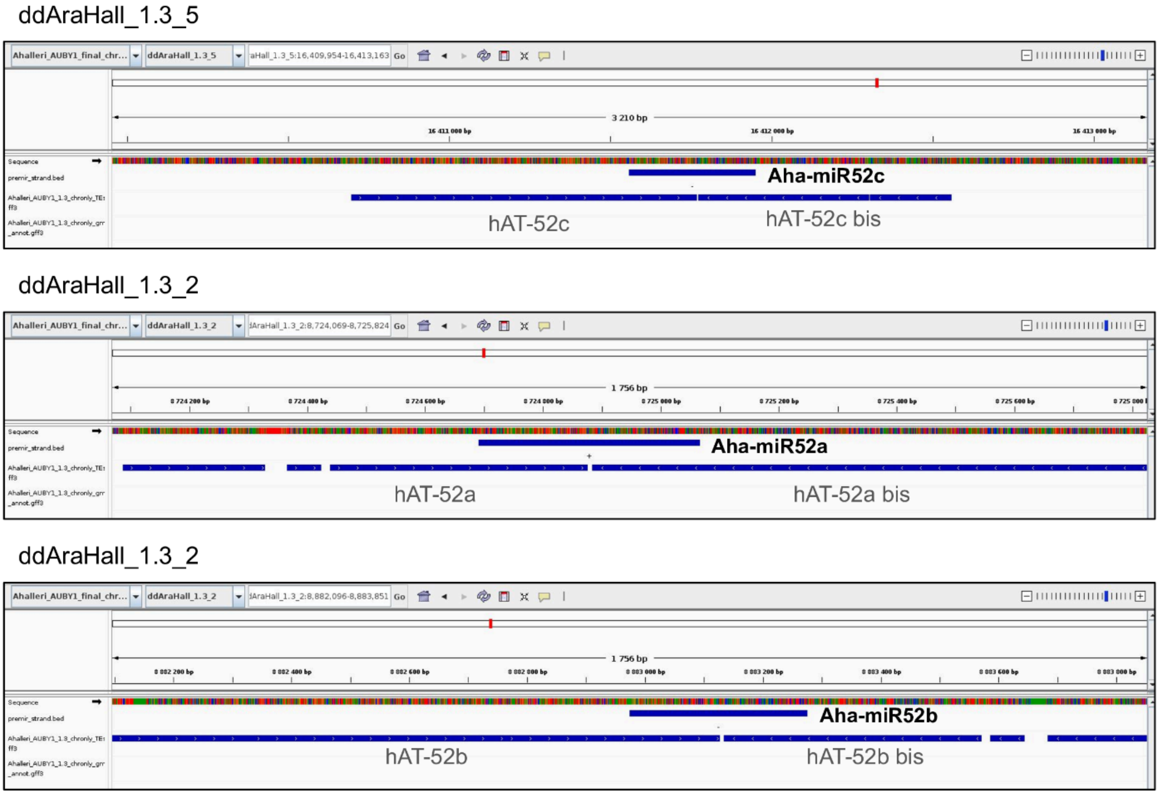


**Figure S3: Aha-miR52 precursor sequences overlap hAT transposons.** Genomic localizations of Aha-miR52a, Aha-miR52b and Aha-miR52c and hAT transposons are visualized with IGV v.2.13.0 (Robinson et al., 2011).

**Duplication class** : direct duplication of a locus containing a previous intralocus inverted duplication

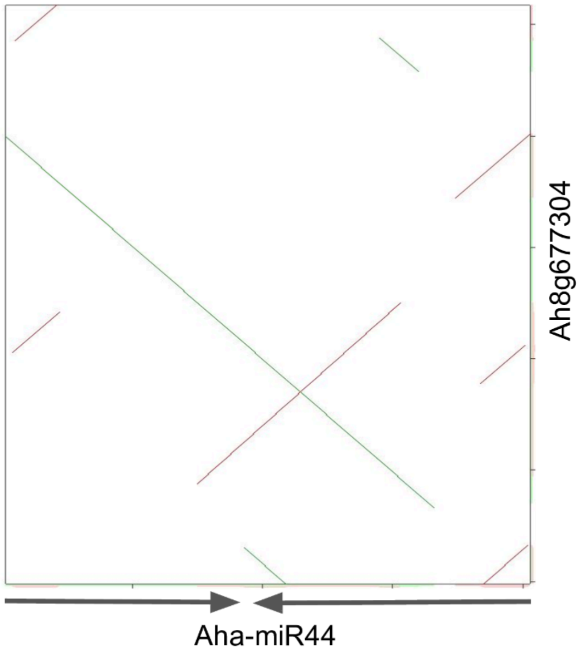


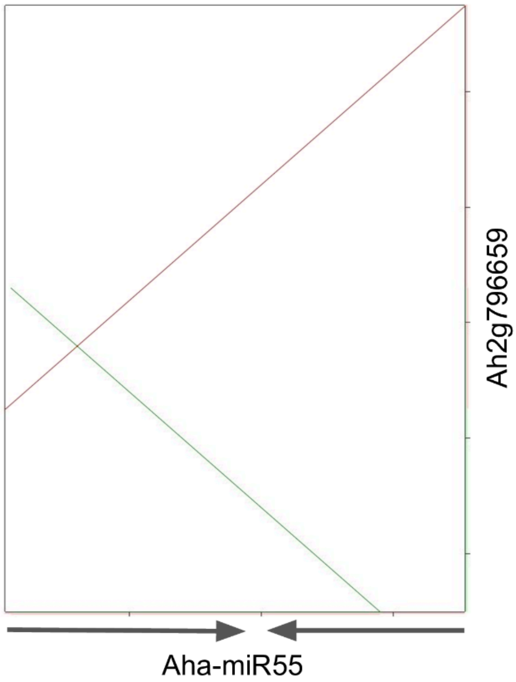


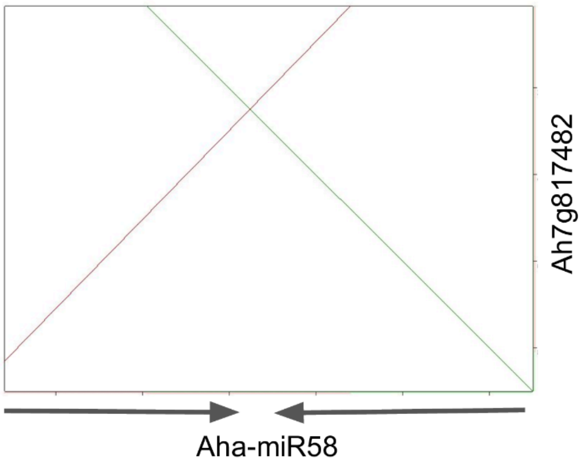


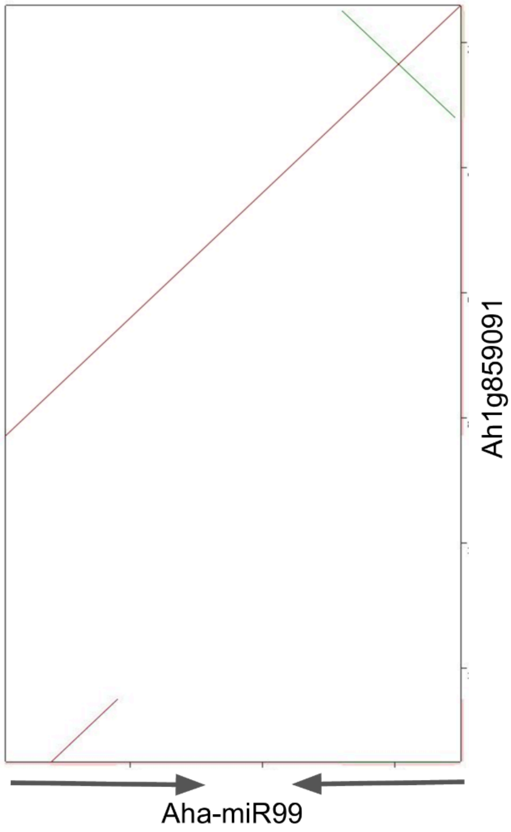

**Duplication class** : inverted duplication of a locus

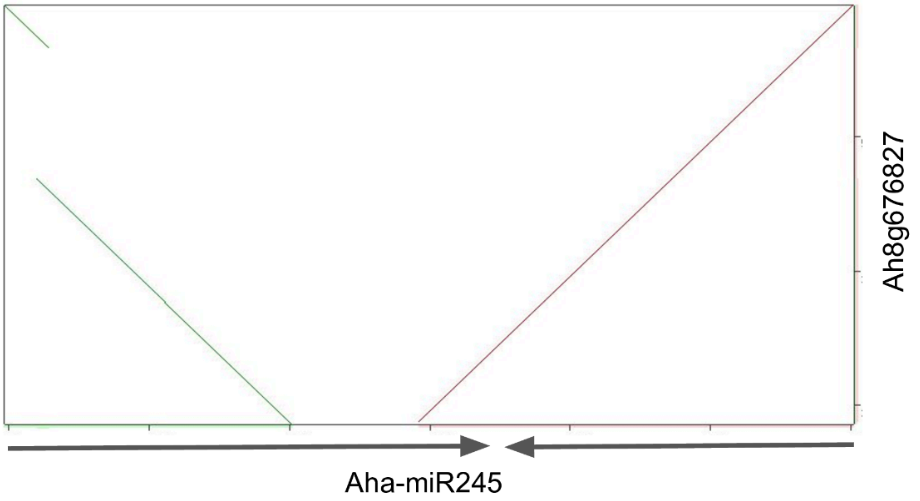


**Figure S4: Comparison between the structure of the miRNA precursor and the portion of the CDS to which it has sequence similarity.** The miRNA precursors can originate from a direct duplication of a portion of the protein-coding gene already containing an intra-CDS inverted duplication, or from an inverted duplication of a portion of a CDS. The dotplots were produced by YASS (Noe and Kucherov, 2005). The green lines represent direct repeats, while the red ones represent inverted repeats. The stem-loop structure of the miRNA precursors is represented by the two inverted arrows, the loop being located between the two arrows.
